# Effects of Perinatal Fluoxetine Exposure on Novelty-induced Social and Non-Social Investigation Behaviors in a Seminatural Environment

**DOI:** 10.1101/2021.02.09.430401

**Authors:** Ole Christian Sylte, Jesper Solheim Johansen, Indrek Heinla, Danielle J Houwing, Jocelien DA Olivier, Roy Heijkoop, Eelke MS Snoeren

## Abstract

Selective serotonin reuptake inhibitors (SSRIs) are increasingly prescribed as medication for various affective disorders during pregnancy. SSRIs cross the placenta and affect serotonergic neurotransmission in the fetus, but the neurobehavioral consequences for the offspring remain largely unclear. Recent rodent research has linked perinatal SSRI exposure to alterations in both social and non-social aspects of behavior. However, this research has mainly focused on behavior within simplified environments. The current study investigates the effects of perinatal SSRI exposure on social and non-social investigation behaviors of adult rat offspring upon introduction to a novel seminatural environment with unknown conspecifics. During the perinatal period (gestational day 1 until postnatal day 21), rat dams received daily treatment with either an SSRI (fluoxetine, 10 mg/kg) or vehicle. Adult male and female offspring were observed within the first hour after introduction to a seminatural environment. The results showed that perinatal fluoxetine exposure altered aspects of non-social investigation behaviors, while not altering social investigation behaviors. More specifically both fluoxetine exposed males and females spent more total time on locomotor activity than controls. Furthermore, fluoxetine exposed females spent less time exploring objects and specific elements in the environment. The data suggest that perinatal exposure to SSRIs leads to a quicker, less detailed investigation strategy in novel environments, and that the alteration is mostly pronounced in females.

## 1. Introduction

A considerable number of women experience depression or other mental disorders during pregnancy. Approximately 1 in 10 pregnant women fulfill the DSM-5 diagnostic criteria for major depressive disorder (Bennett et al. 2004; Woody et al. 2017). In treatment of maternal depression and anxiety, selective serotonin reuptake inhibitors (SSRIs) are the most frequently prescribed class of drugs, as it has been considered relatively safe for both mother and child. The prescription rate of SSRIs to pregnant women has increased tremendously in the last decades (Mitchell et al. 2011), and recent estimates suggest a worldwide prevalence of 3% (Molenaar et al. 2020) with significant geographically differences (Andrade et al. 2008; Charlton et al. 2015). Consequently, hundreds of thousands of babies exposed to SSRIs during early development are born every year. Despite the widespread use, we have limited knowledge on whether SSRI exposure during the early stages of brain development can lead to altered long-term behavioral outcomes, such as social and non-social behaviors.

Antidepressants, such as SSRIs, reach the fetus by crossing the placenta and are present in breast milk (Kristensen et al. 1999; Rampono et al. 2004). Thus, children can potentially be exposed to SSRIs during the entire perinatal period (Kim et al. 2006; Noorlander et al. 2008). SSRIs inhibit the function of the serotonin-reuptake transporter (SERT or 5-HTT), which leads to an accumulation of 5-HT in the synaptic cleft. This in turn increases the magnitude and duration of 5-HT activity at pre- and post-synaptic 5-HT receptors. In the adult brain, 5-HT acts mainly as a modulatory neurotransmitter, regulating emotion, cognition, sleep and stress responses (Olivier et al. 2011a). However, in the developing brain, 5-HT is widespread and acts as a neurotrophic factor regulating cell division, differentiation, migration, and synaptogenesis (Azmitia 2001; Gaspar et al. 2003). Consequently, developmental SSRI exposure is suggested to affect both neurodevelopment and later-life behaviors (Muller et al. 2016).

Previous studies in humans have shown associations between developmental SSRI exposure and impaired social behavior (Klinger et al. 2011), increased risk of speech and language disorders (Brown et al. 2016), and elevated levels of internalizing behavior, like anxiety and depression (Hermansen et al. 2016; Lupattelli et al. 2018; Malm et al. 2016), as well as increased risk of attention-deficit hyperactivity disorders (Man et al., 2018). While the existing literature has mainly examined the childhood years, little is known on whether these associations persist into adulthood. In addition, outcomes such as depression may not emerge before a certain age and could therefore remain undiscovered.

Epidemiological research on humans, like the above-mentioned studies, are correlational in nature, and do not necessarily imply causation. A frequent problem with human studies is the difficulty to isolate the effects of SSRI exposure from the effects of maternal mental health. Women using SSRIs during pregnancy are likely suffering from depression, which itself has been shown to have negative impact on the offspring (Dunkel Schetter 2011; El Marroun et al. 2014; Goodman 2007). Animal research, on the other hand, allows to control for potential interference from confounding factors, like maternal health, drug dose and timing of exposure. As rodent and human serotonergic development is remarkably similar (Glover and Clinton 2016), rodent studies can provide valuable translational insight about how developmental SSRI exposure affects human offspring.

Rodent studies investigating the effects of developmental exposure to SSRIs have reported alterations in different social and non-social behaviors in the offspring. In juvenile male and female offspring, both pre- and post-natal SSRI exposure have been shown to decrease social play behavior (Houwing et al. 2019b; Khatri et al. 2014; Olivier et al. 2011b; Rodriguez-Porcel et al. 2011; Simpson et al. 2011). Similar tendencies were found in adult rats with developmental SSRI exposure leading to less social interactions (Olivier et al. 2011b; Rodriguez-Porcel et al. 2011), or decreased interest to explore a novel conspecific (Khatri et al. 2014; Rodriguez-Porcel et al. 2011; Simpson et al. 2011; Zimmerberg and Germeyan 2015). SSRI exposure can also decrease (Houwing et al. 2020), or increase (Gemmel et al. 2017; Kiryanova and Dyck 2014; Svirsky et al. 2016), aggressive-like social behaviors. Furthermore, a recent meta-analysis revealed that developmental exposure to SSRI was linked to reduced activity and explorative behaviors in adult rats and mice (Ramsteijn et al. 2020).

Most rodent studies, however, have used simplified test set-ups which only investigates a small fraction of all behaviors. Furthermore, these studies do not account for the environmental and social complexity of real-world situations. To bypass these limitations, recent studies from our research group have employed a seminatural environment enabling rats to express many aspects of their natural behaviors (Hegstad et al. 2020; Heinla et al. 2020; Houwing et al. 2019a). These studies showed that perinatal SSRI fluoxetine (FLX) exposure leads to various alterations in social and non-social behaviors in a naturalistic setting. More specifically, perinatal fluoxetine exposure was associated with an increased amount of passive social behaviors in both males and females, but a reduction of active social behavior, general activity (Houwing et al. 2019a), and pro-social behaviors in females (Heinla et al. 2020). Interestingly, these studies were performed in the seminatural environment *after* the rats were familiarized to each other and the physical environment. It is currently unknown how social and non-social behaviors manifest directly after introduction to a novel environment with unfamiliar conspecifics. As perinatal SSRI exposure seem to alter stress-coping behaviors (Houwing et al. 2019a), one could hypothesize that the stressor of a novel environment with new conspecifics could lead to more pronounced changes in social and non-social behaviors.

The aim of the current study was to investigate if perinatal SSRI exposure alters social and non-social investigation behaviors in a novel environment with unknown conspecifics. We define investigation as behaviors that provides the animal with information about a novel stimulus. More specifically, social investigation refers to when the stimulus investigated is a conspecific, such as when sniffing and grooming others, while non-social investigation refers to investigation of inanimate objects and environmental locations. In line with previous studies (Heinla et al. 2020; Houwing et al. 2019a), we expected perinatal fluoxetine exposure to show a reduction in active social behavior in non-social investigation (exploratory) behavior in the initial phase of the introduction to the seminatural environment. In addition, as introduction to a new environment can be considered a stressful situation, we also expected to observe an increase in self-grooming behavior in FLX-exposed animals.

## 2. Material and Methods

The data was collected from video recordings obtained in a previously performed experiment (Houwing et al. 2019a). The materials and methods are therefore similar to our previous studies using data from the same experiment (Hegstad et al. 2020; Heinla et al. 2020; Houwing et al. 2019a). However, the behavioral scoring scheme and the time window of observation was unique for the current study.

### 2.1 Animals and dam housing

A total of 20 Wistar rats (10 males, 10 females), weighing 200-250 grams on arrival, were obtained from Charles River (Sulzfeld, Germany) for breeding. After arrival, same-sex pairs were housed in Makrolon IV cages (60 x 38 x 20 cm) on a reversed 12:12 hours light/dark cycle, in which the lights were turned on at 23.00. Temperature in the room was 21 ± 1°C, and the relative humidity was 55 ± 10 %. Standard rodent food pellets (standard chow, Special Diets Services, Witham, Essex, UK), water and nesting material were available ad libitum. Animal care and experimental procedures were conducted in agreement with European Union council directive 2010/63/EU. The protocol was approved by the National Animal Research Authority in Norway.

### 2.2 Breeding and antidepressant treatment

Daily, all females were checked for sexual receptivity by placing them together with a male rat for 5 minutes. When lordosis behavior was observed, they were considered in proestrus and thus ready for breeding. The female then got placed together with a male in an isolated Makrolon IV cage for the next 24 hours (gestational day 0). Afterwards, they returned to their initial same-sex pairs for the first two weeks of pregnancy. From gestational day 14, the females were placed solitarily until delivery (gestational day 21/postnatal day 0).

During the 6-week period from conception (gestational day 0) to weaning (postnatal day 21), females received either the SSRI fluoxetine 10 mg/kg (Apotekproduksjon, Oslo, Norway) or vehicle (methylcellulose; Sigma, St. Louis, MO, USA) daily by oral gavage. The offspring were thus exposed to perinatal fluoxetine via the treatment of the dams (in utero and via breast feeding). The fluoxetine treatment was prepared with tablets for human usage that were pulverized and dissolved in sterile water (2mg/mL) and injected at a volume of 5mL/kg. Methylcellulose powder, the non-active filling of a fluoxetine tablet, was used as control condition. The powder was dissolved in sterile water to create a 1% solution and administered at a volume of 5mL/kg as well. Every third day, females were weighed to ensure correct dosage of fluoxetine/vehicle. The chosen dosage of fluoxetine was decided upon comparison of fluoxetine blood levels of humans and animals (Lundmark et al. 2001; Olivier et al. 2011b). When the rat dams got close to the end of pregnancy, they were checked two times a day (09.00 and 15.00) for delivery.

### 2.3 Offspring housing

The offspring were housed together with their mothers until weaning (gestational day 21). After weaning, groups of two or three same-sex littermates were housed together in Makrolon IV cages (see cage distribution in the supplemental materials, Table S1). They were left undisturbed, except for the ovariectomy (see section for Procedure) and weekly cage cleaning, until introduction to the seminatural environment at the age of 13-18 weeks. To enable individual recognition, ears were punched. In Figure 1, a schematic overview shows all experimental procedures from gestational day 0 to the end of the experiment.

**Figure 1.**
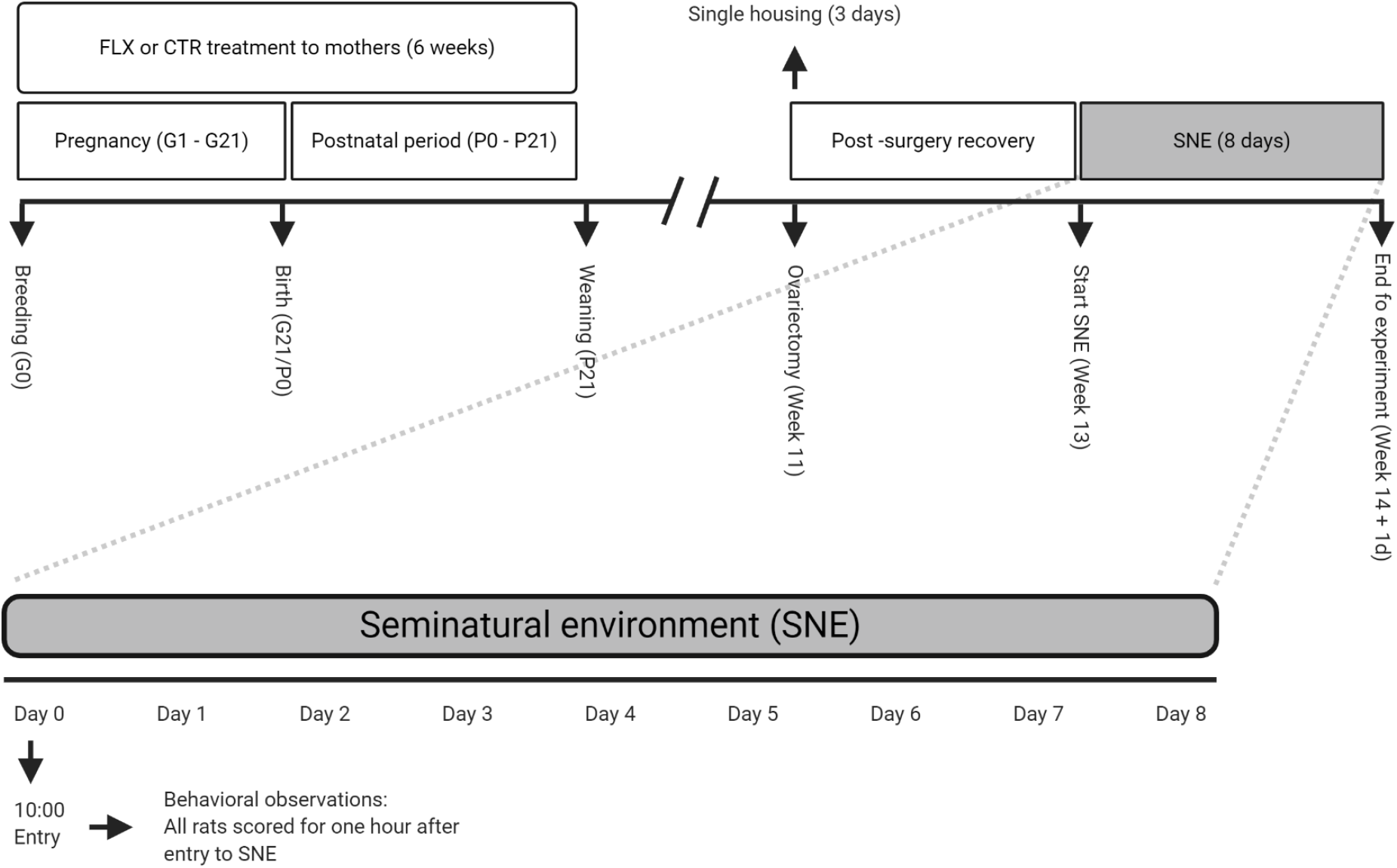
Overview of experimental procedures. FLX = fluoxetine, CTR = control, P = postnatal day, G = gestational day. Created with BioRender (https://biorender.com/).

### 2.4 Seminatural environment

The seminatural environment (SNE; 240 x 210 x 75 cm) consisted of two parts: an open area and a burrow system (Figure 2; (Chu and Agmo 2014; Houwing et al. 2019a; Snoeren et al. 2015)). Four openings (8 x 8 cm) connected the two areas. In the open area, two partitions (40 x 75 cm) simulated natural obstacles. The burrow system consisted of connected tunnels (width 7.6 cm, height 8 cm) and four nest boxes (20 x 20 x 20 cm). Plexiglas covered the burrow at the height of 75 cm, while the open area remained open. A curtain between the two parts allowed for different light settings. The burrow was left dark the entire time. In the open area, on the other hand, light settings simulated a day-night cycle. A lamp located 2,5 m above the floor, simulated daylight (180 lux) between 22.45 and 10.30. From 10.30 to 11.00 the lights gradually decreased to 1 lux (simulating moonlight). The darkness lasted until the light gradually increased from 1 to 180 lux between 22.15 and 22.45.

**Figure 2.**
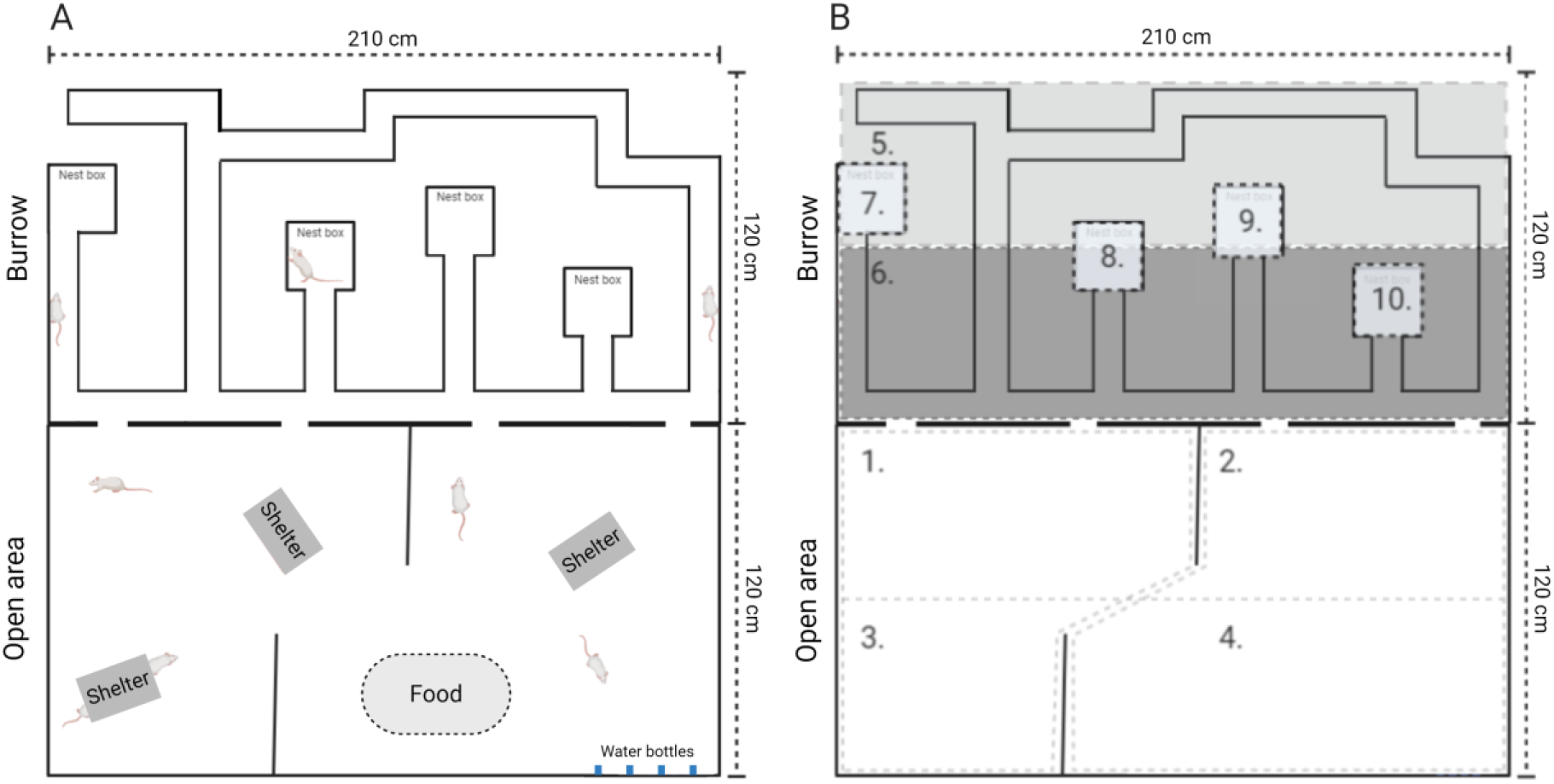
The Seminatural Environment. Illustration of the whole seminatural environment (A) and sectioning of the different locations (B). 1 = open area close to burrow left, 2 = open area close to burrow right, 3 = open area far away from burrow left, 4 = open area far away from burrow right, 5 = tunnels far away from open a, 6 = tunnels close to OA, 7 = nestbox left, 8 = nestbox mid-left, 9 = nestbox mid-right, 10 = nestbox right. Created with BioRender (https://biorender.com/). A picture of the seminatural environment can be found in the Supplemental materials (Figure S1).

The whole ground of the SNE was covered with a layer (2 cm) of aspen wood chip bedding (Tapvei, Harjumaa, Estonia). The nest boxes had 6 squares of nesting material in each (non-woven hemp fibers, 5 x 5 fibers, 5 mm thickness, Datesend, Manchester, UK). Three plastic shelters (15 x 16.5 x 8.5 cm, Datesend, Manchester, UK) were placed in the open area. Additionally, 12 aspen wooden sticks (2 x 2 x 10 cm, Tapvei, Harjumaa, Estonia) were randomly placed around in the SNE. A pile of food pellets (approx. 2 kg) and four bottles of water were available at all time (see location in Figure 2A).

Two video cameras (Basler) were mounted on the ceiling, 2 m above the open area (regular camera) and the burrow system (infrared camera) respectively. Media Recorder 2.5 was employed for video recordings. The data got immediately stored on an external hard drive. The recording was manually stopped and restarted every 24 hours. The purpose was to ensure that eventual errors only would affect one day of recorded data.

### 2.5 Design of the study

Initially, five cohorts, each consisting of eight rat offspring, were placed one at the time in the SNE. However, one day of video material was lost due to recording error, which reduced the number of cohorts to four. A cohort consisted of 4 males and 4 females of which each sex constituted 2 controls (CTR) and 2 fluoxetine (FLX) rats. Thus, data from this experiment came from 8 CTR-males, 8 CTR-females, 8 FLX-males and 8 FLX-females (see Table S2 for more details). Within a cohort, same sex rats came from different litters and were thus unfamiliar to each other. Some rats had one sibling from the opposite sex in the same cohort. However, these rats had been housed in different home cages since weaning. Analysis of the data also revealed no differences in behavior towards males or females, suggesting that the sibling factor did not influence the data.

### 2.6 Procedure

For the purpose of a previous study (Houwing et al. 2019a), the female offspring were ovariectomized two weeks before entering the SNE in order to control their estrous cycle. Although irrelevant for the objective of the current study, this procedure had the effect of keeping the females in diestrus of the menstrual cycle during the observation period. Before entering the SNE, the rats were shaved on the back and tail-marked under isoflurane anesthesia for individual recognition (for more details, see (Houwing et al. 2019a)). All rats were also weighed, confirming that there was no weight difference between CTR- and FLX-rats.

Each cohort was placed in the SNE for 8 days. See Figure 1 for an overview of the whole procedure. The cohorts were introduced to SNE on the first day (day 0) at 10.00 and removed on day 8 at the same time. However, only data from the first hour was used for the purpose of this study. All rats were again weighed after being removed from the SNE. No difference in weight was observed between CTR- and FLX-rats. In order to remove olfactory cues, the SNE was cleaned and bedding changed between cohorts.

### 2.7 Behavioral observations

The frequency and/or duration of several behaviors (see Table 1) were scored manually using The Observer XT, version 12 (Noldus, Wageningen, The Netherlands). Two observers, blinded for the animal treatment, independently scored either males or females across all four cohorts. In addition to behavior, (1) location of the animal (see Figure 2B), (2) whether the animal initiated the respective behavior or was respondent to it, (3) whether the animal was in physical contact with another animal or not during the respective behavior, and lastly, (4) ID of the interacting partner was scored. Since we were interested in observing how the rats behaved in a novel environment with unfamiliar conspecifics, all rats were scored in the first 60 minutes after entry to the SNE.

**Table 1.** Description of recorded behaviors.

| Behavior | Description |
| --- | --- |
| Walking/running | Walking or running through the environment |
| Chasing | Running forward in the direction of a conspecific |
| Non-social exploration | Exploring the environment by sniffing, usually when slowly walking or sitting still |
| Digging | Digging, pushing or carrying bedding/nesting/food material |
| Resting/immobile alone | Sitting or sleeping with minimal movement of the head without other rats in close vicinity |
| Resting/immobile socially | Sitting or sleeping with minimal movement of the head with at least 1 other rat on maximum 1 rat body length away |
| Hiding alone | Being in the shelter alone |
| Hiding socially | Being in the shelter with at least one other rat |
| Following | Walking or running in the same direction as another rat in front. |
| Allogrooming | Grooming any part of a conspecific's body, usually on the head or in the neck region |
| Sniffing anogenitally | Sniffing the anogenital region of the conspecific |
| Sniffing nose-to-nose | Sniffing the facial region of the conspecific |
| Sniffing body | Sniffing any part of the conspecifics body, except for the anogenital and facial region |
| Fighting | Kicking, pouncing, pushing, grappling, boxing or wrestling another rat |
| Nose-off | Facing another rat, usually in a tunnel, resulting in one rat moving forward and the other backing up |
| Self-grooming | Grooming itself |
| Freezing | Complete absence of movement in addition to a tense body posture |
| Rearing supported | Raising itself upright on its hind paws, facing a wall or an object |
| Rearing unsupported | Raising itself upright on its hind paws, not facing a wall or an object |

### 2.8 Data preparation and statistical analysis

As shown in Table 2, the recorded behaviors were combined into behavioral clusters. For each rat, we calculated the total duration and the number of events for every behavior and behavioral cluster. This data was later divided into six 10-minute time-bins in order to analyze behavioral changes over time. Latencies to meet the other rats, and latencies to visit the different locations of the SNE were also noted. This data was later divided and analyzed cumulative over the first 1, 3, 5, 10, 20, 30, and 60 minutes. In this study, we operationalized social investigation behaviors as the cluster “socially active behaviors” and the latencies to meet all other rats, whereas non-social investigation behaviors were operationalized as the cluster “general activity” and latencies to visit all the locations (See Figure 2B).

**Table 2.** Description of behavioral clusters.

| Cluster | Behaviors within clusters |
| --- | --- |
| Socially active behaviors | Sniffing anogenitally, sniffing nose-to-nose, sniffing body, and allogrooming |
| General activity | Walking/running, non-social exploration |
| Non-socially passive behaviors | Resting alone, hiding alone |
| Socially passive behaviors | Hiding socially, resting socially |
| Conflict behaviors | Nose-off, fighting |

Normality of data was determined with Shapiro-Wilks tests. Data with p < .05 was analyzed non-parametrically. Simple group comparisons were performed with either a student t-test or the non-parametric Mann-Whitney U test. Repeated measures ANOVA was used when the behaviors were analyzed over time. In cases the Mauchly’s test indicated violation of sphericity from the ANOVA output, the degrees of freedom were corrected using Greenhouse-Geisser estimates of sphericity. To correct for multiple comparisons, the Benjamini-Hochberg procedure was performed on all significant results together with a predetermined set of variables (sniffing, self-grooming, non-social exploration, conflict behaviors). All tests reported were done 2-tailed.

Because male and female behavior were scored by two different observers, no conclusions were drawn regarding potential differences between males and females.

### 2.9 Statement Open Science Framework (OSF)

The design of our study was preregistered on OSF on the 25th of March 2019 (https://osf.io/m87j5). There were no changes in analysis, except that we did not use the originally planned additional control group. As stated at OSF, the planned control group was not suitable, because it consisted of aged rats and had a different composition in number of rats (7 versus 8). We therefore concluded that these differences would make it impossible to compare the cohorts of the current study.

## 3 Results

From the behavioral scoring, we obtained a lot of data. A complete overview of all behaviors can be found in Table S3 and S4. In this result section, we only mention the relevant behavior to the purpose of the paper. A summary of the main finding are provided in Table 3.

**Table 3.**
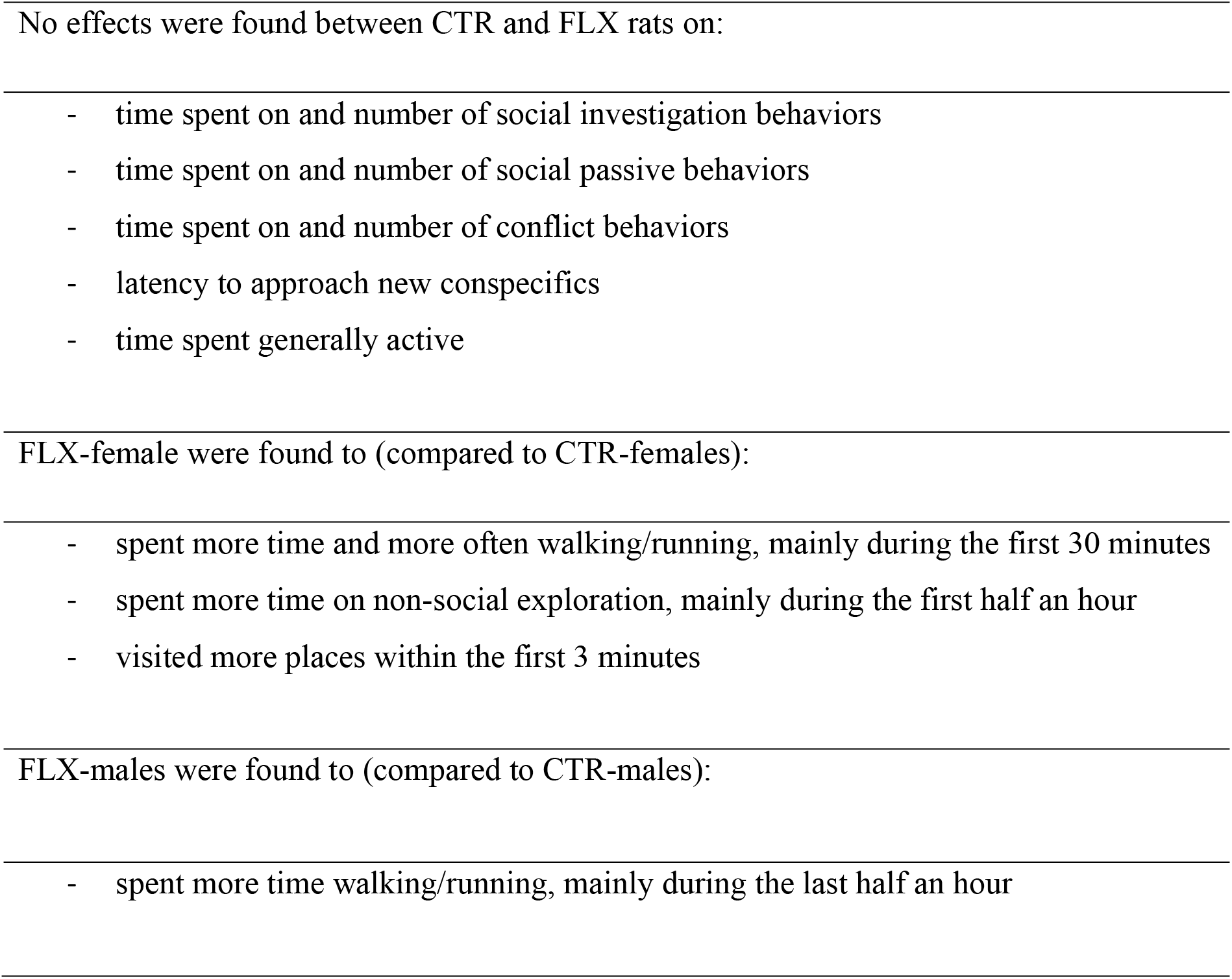
Summary of main findings.

### 3.1 Social investigation behaviors

The data analysis revealed that CTR- and FLX-females did not differ in time spent on (*t* = −1.04, *p* = .315, *d* = −0.52, Figure 3A) or number of episodes (*t* = −1.04, *p* = .318, *d* = −0.52) performing socially active behaviors. When looking separately at the different behavioral components constituting the cluster (see Table 2), CTR- and FLX-rats did not differ on any other behavioral components constituting the clusters relevant to social behaviors (socially active behaviors, socially passive behaviors and conflict behaviors). No difference was found between CTR- and FLX-males for socially active behaviors in total time (*t* = 0.95, *p* = .356, *d* = 0.48, Figure 3D) or on number of episodes (*t* = 0.103, *p* = .919, *d* = 0.05).

**Figure 3.**
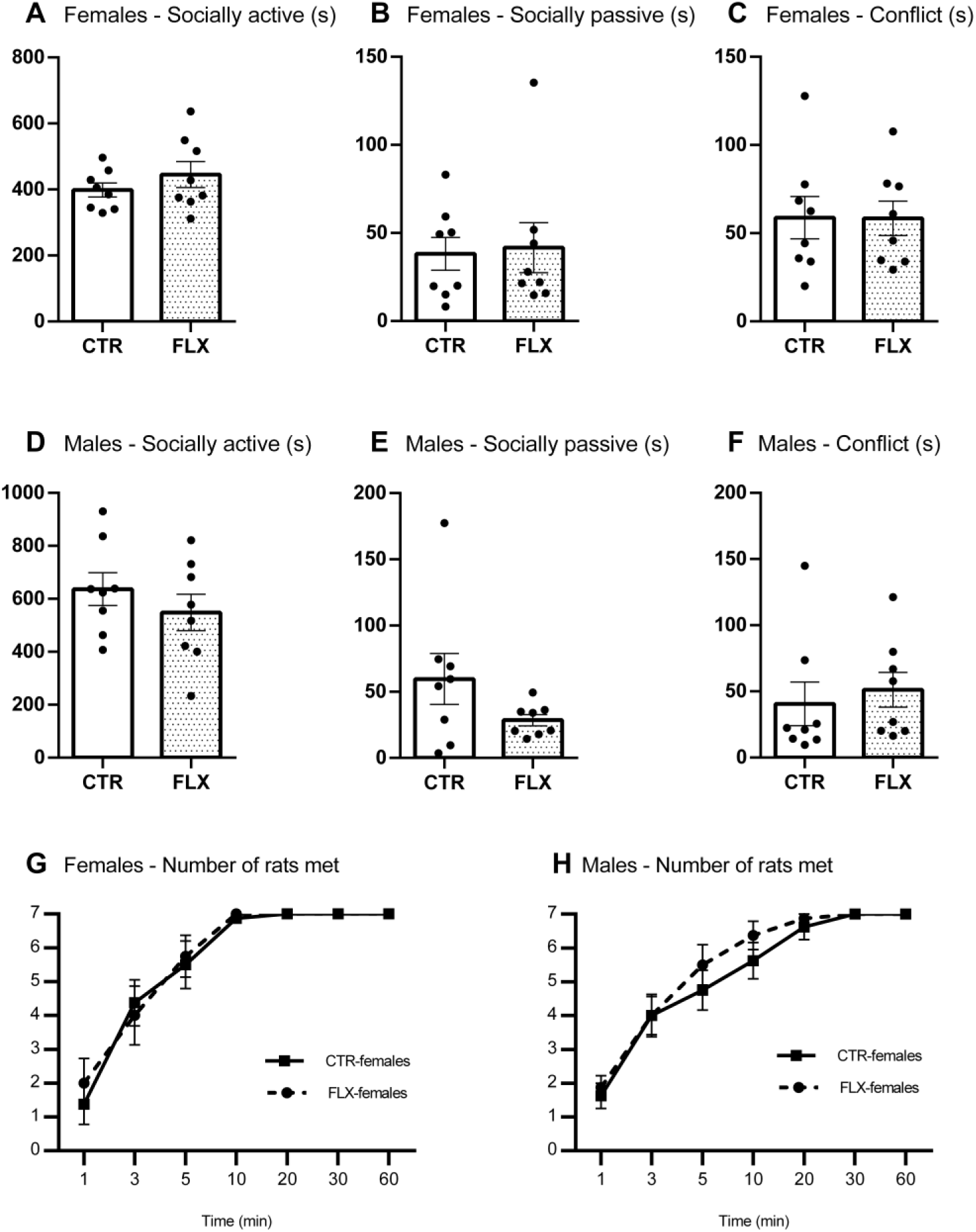
Social behaviors in females and males. The data represent the time spent (s) on socially active behaviors (A, D), socially passive behaviors (B, E), conflict behaviors (C, F), and the total number of rats met over time (G, H). All graphs show comparisons between CTR-females (n = 8) and FLX-females (n = 8) or between CTR-males (n = 8) and FLX-males (n = 8). Data are shown with individual data points with bars representing the group means (A-F), or with squares and circles representing respective group means (G-H). Error bars are representing SEM.

Although the treatment groups did not differ in the amount of socially active behaviors, it could still be the case that the groups had different interest in meeting other rats. To investigate this possibility, we first looked at the latencies to when the rats had met all seven other cohort-members. The data analysis showed that there was no significant difference in latency to meet all cohort-members between CTR- and FLX-rats for females (*t* = 0.84, *p* = .418, *d* = 0.42) or males (*U* = 24.00, *z* = −0.84, *p* = .422, *r* = −.21). We subsequently measured how many cohort members the rats had met as a function of time. CTR- and FLX-rats were compared on cumulative data measured at 1/3/5/10/20/30/60 minutes. For FLX-females, there were no significant differences in the number of rats met (treatment effect: *F*(1,14) = 0.05, *p* = .821) or in the pattern of rats met (timepoints x treatment: *F*(1.73, 24.24) = 0.28, *p* = .725) over time compared to CTR-females (Figure 3G). Similarly, CTR- and FLX-males did not differ in the number of rats met across all timepoints (treatment effect: *F*(1,14) = 0.49, *p* = .492) or in the pattern of rats met over time (timepoints x treatment: *F*(2.05, 28.74) = 0.59, *p* = .563, Figure 3H).

### 3.2 Other social behaviors

We also investigated some other social behaviors, such as socially passive behaviors and conflict behaviors. No difference was found between CTR- and FLX-females in total time (*U* = 33.00, *z* = 1.05, *p* = 1, *r* = .03, Figure 3B) or number of episodes being socially passive (*t* = −0.28, *p* = .784, *d* = −0.14). Furthermore, CTR- and FLX-females spent a similar amount of time (*t* = 0.03, *p* = .978, *d* = 0.01, Figure 3C) and episodes (*t* = −0.40, *p* = .692, *d* = −0.20) in conflict with other rats. Similarly, for males, no differences were found for time spent on social passive behavior (*U* = 41.00, *z* = 0.95, *p* = .382, *r* = .24, Figure 3E), episodes of social passive behavior (*t* = 1.48, *p* = .161, *d* = 0.74), time spent on conflict behavior (*t* = −0.03, *p* = .655, *d* = −0.02, Figure 3F), or episodes in conflict behavior (*U* = 42.00, *z* = 1.05 *p* = .786, *r* = .26).

### 3.3 Non-social investigation behaviors

CTR- and FLX-females did not differ in time spent on (*t* = −1.04, *p* = .311, *d* = 0.31, Figure 4A) or in the number of episodes of general activity (*t* = −1.82, *p* = .090, *d* = −.0.91). However, FLX-females were found to spend significantly *more* time walking/running (*U* = 56.00, *z* = 2.52, *p* = .025, *r* = .63, Figure 4B) but *less* time on non-social exploration (*U* = 8.00, *z* = −2.52, *p* = .025, *r* = − .63, Figure 4C) compared to CTR-females. FLX-females were also found to have more episodes of walking/running compared to CTR-females (*t* = −4.29, *p* = .005, *d* = −2.15). CTR- and FLX-females did not differ in the number of non-social exploration episodes (*t* = −0.54, *p* = .693, *d* = −0.27). Similar as for the females, no difference in time spent on (*t* = −1.69, *p* = .114, *d* = −0.85, Figure 4D) or on number of episodes in general activity (*t* = −1.60, *p* = 0.131, *d* = −0.80) were found between CTR- and FLX-males. However, just as FLX-females, FLX-males spent more time walking/running than CTR-males (*t* = −3.05, *p* = .045, *d* = −1.52, Figure 4E), but there was no difference in time spent on non-social exploration (*t* = 0.06, *p* = .953, *d* = 0.03, Figure 4F). FLX-males did not differ from CTR-males in the number of episodes walking/running (*t* = −1.61, *p* = .130, *d* = −0.80) or non-social exploration (*t* = −0.73, *p* = .786, *d* = −0.36).

**Figure 4.**
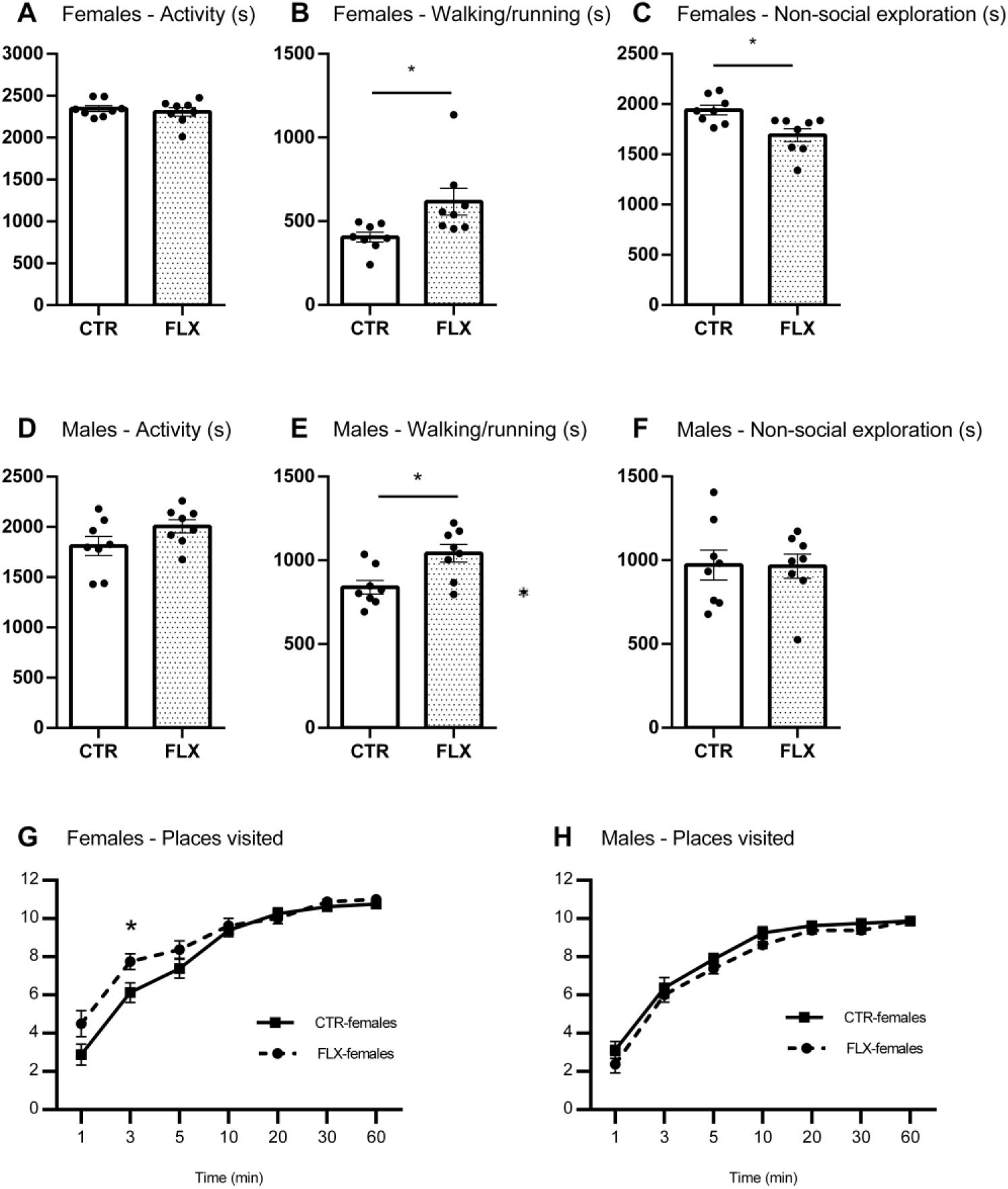
Non-social behaviors in females and males. The data represent the time spent (s) on general activity (A, D), walking/running (B, E), non-social exploration (C, F), and the total number of places in the SNE visited over time (G, H). All graphs show comparisons between CTR-females (n = 8) and FLX-females (n = 8) or between CTR-males (n = 8) and FLX-males (n = 8). Data are shown with individual data points with bars representing the group means (A-F), or with squares and circles representing respective group means (G-H). Error bars are representing SEM. *p < 0.05

We then investigated whether there were differences between CTR- and FLX-rats in how long it took them to visit all the 10 predefined locations (see Figure 2B) of the seminatural environment. Rats that did not visit all locations within the observation time were given a score of 3600 (total observation time in seconds). The results indicated that FLX-rats, both males and females, did not need significantly more or less time to visit all locations than CTR-rats (females: *t* = 1.33 = *p* = .212, *d* = 0.42; males: *t* = −1.15, *p* = .271, *d* = −0.57). We thereafter investigated how many locations the rats visited as a function of time (1/3/5/10/20/30/60 minutes), measured on cumulative data. FLX-females were not significantly faster at visiting the different locations compared to CTR-females (Figure 4G), but when the different time-points were analyzed separately, they seem to have visited significantly more locations within the first 3 minutes (*t* = −2.46, *p* = .027, *d* = −1.23) compared to CTR-females. No difference in the number of locations visited (treatment effect: *F*(1,14) = 3.43, *p* = .085) or in the pattern (time x treatment: *F*(2.64, 36.97) = 0.39, *p* = .735) over time were found between the CTR- and FLX-males (Figure 4H).

### 3.4 Other non-social behaviors

We also looked at other relevant non-social behaviors, including non-socially passive behaviors. The analysis revealed that there was no significant difference between CTR- and FLX-females in time spent on (*U* = 28.00, *z* = −0.42, *p* = .721, *r* = −0.11) or in the number of non-socially passive behaviors (*t* = −0.12, *p* = .903, *d* = −0.06). Similarly, for the male groups, no significant difference was found for time spent on (*t* = 1.62, *p* = .127, *d* = 0.81) or in the number of non-socially passive behaviors (*t* = 0.62, *p* = .546, *d* = 0.31)

Next, we investigated whether CTR- and FLX-rats showed different level of anxiety/stress-related behaviors. The results revealed no significant difference between CTR- and FLX-rats for time spent on (females: *t* = 1.67, *p* = .195, *d* = 0.84; males: *U* = 37.00, *z* = 0.53, *p* = .806, *r* = .13) or in the number of episodes (females: *t* = 0.58, *p* = .693, *d* = 0.29; males: *t* = −0.60, *p* = .860, *d* = −0.30) self-grooming. When investigating the total time in the open area, no significant difference was found between CTR- and FLX-rats (females: *t* = − 1.39, *p* = .186, *d* = −0.70; males: *t* = −0.98, *p* = .345, *d* = −0.49). Similarly, the treatment groups did not differ on the total time spent in the burrow area (females: *t* = 1.57, *p* = .138, *d* = 0.79; males: *t* = 1.02, *p* = .323, *d* = 0.51).

### 3.5 Behavioral adaption over time

Finally, we were interested to see whether the treatment groups adapted differently to the novel physical and social environment, and thus, whether the differences in behavior between the groups were stable over time. We therefore divided the dataset into six 10-minutes time-bins and assessed the differences between CTR- and FLX-rats on social and non-social behaviors over the course of the observation period.

The repeated measure analysis revealed that FLX-females and FLX-males did not show a significantly different pattern of time spent on socially active behaviors, compared to CTR-females (time-bin x treatment: *F*(5,70) = 0.26, *p* = .932, η_p_^2^ = .02, Figure 5A) or CTR-males (time-bin x treatment: *F*(5,70) = 0.51, *p* = .765, η_p_^2^ = .04, Figure 5B) respectively. Similarly, when looking at the frequency of socially active behaviors, no interaction between time-bin and treatment was found for female (*F*(5,70) = 0.63, *p* = .675, η_p_^2^ = .04) or male rats (*F*(5,70) = 0.99, *p* = .431, η_p_^2^ = .07).

**Figure 5.**
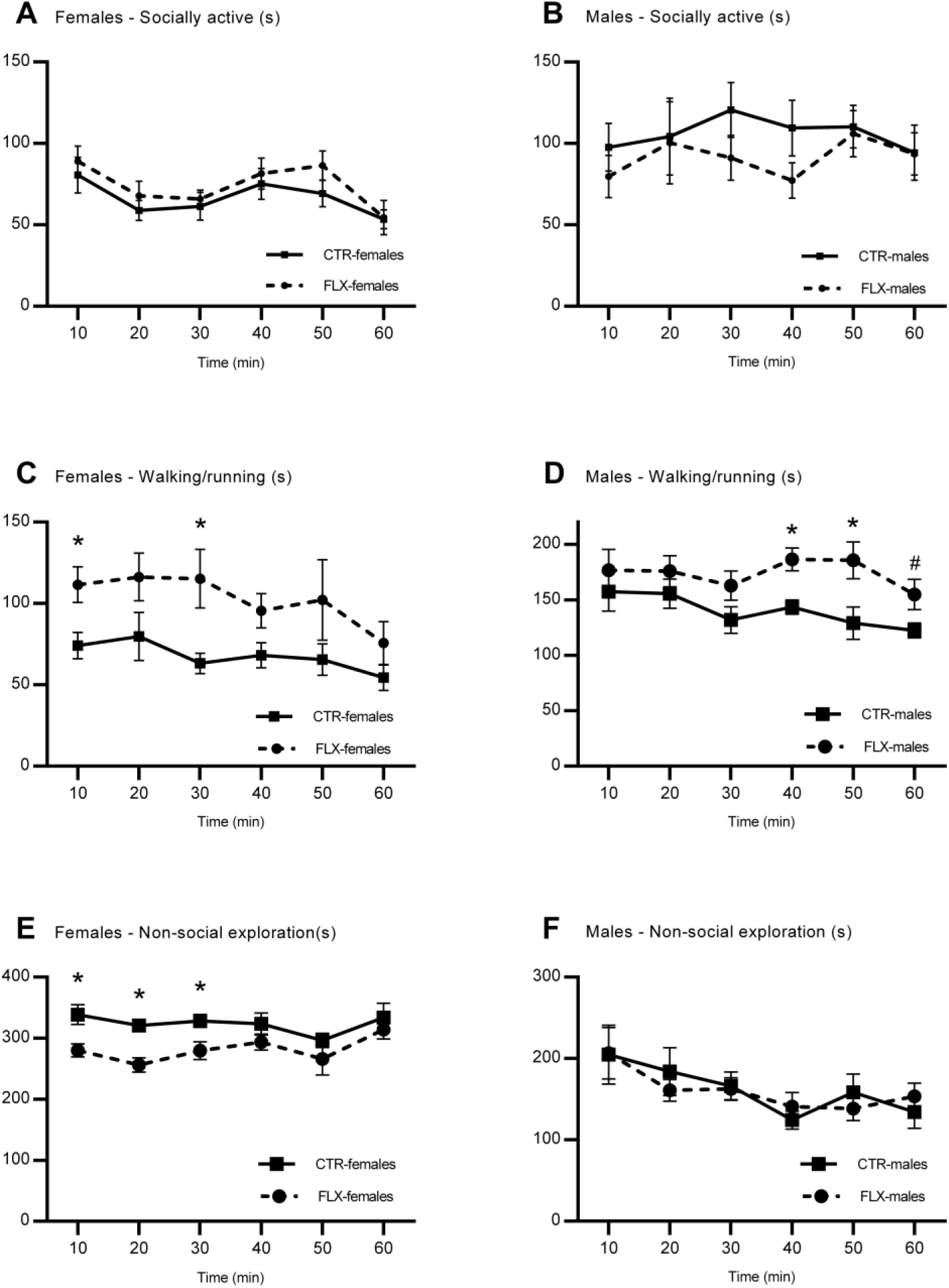
Social and non-social investigation behaviors measured as a function of time. The data represent the time spent (s) on different behaviors as a function of time measured every 10 minutes. The graphs show socially active behaviors (A, B), walking/running (C, D), and non-social exploration (E, F). All graphs show comparisons between CTR-females (n = 8) and FLX-females (n = 8) or between CTR-males (n = 8) and FLX-males (n = 8). Squares and circles represent respective group means, error bars representing ± SEM.*p < 0.05, #p < 0.06

For time spent on walking/running, no differences as a function of time were found between the CTR- and FLX-rats for females (*F*(5,70) = 0.63, *p* = .679, η_p_^2^ = .04) or males (*F*(2.64. 36.92) = 0.69, *p* = .634, η_p_^2^ = .05), meaning that the increase in walking/running was present during the whole course of the hour and was most pronounced during the first 10-(*t* = −2.77, *p* = .015, *d* = −1.38) and 30-minutes (*U* = 59.00, *z* = 2.84, *p* =.003, *r* = .71) in FLX-females, and during the first 40-(*t* = −3.58, *p* = .003, *d* = −1.79) and 50-minutes (*t* = −2.56, *p* = .023, *d* = −1.28) in FLX-males, compared to CTR-animals. Similar results were found when analyzing the frequency of walking/running (females: *F*(5,70) = 0.88, *p* = .498, η_p_^2^ = .06; males: *F*(2.85, 39.88) = 0.82, *p* = .483, η_p_^2^ = .06). In term of non-social exploration, neither FLX-females (*F*(5,70) = 0.84, *p* = .529, η_p_^2^ = .06) nor FLX-males (*F*(2.87, 40.20) = 0.47, *p* = .697, η_p_^2^ = .03) showed a significant different pattern of time spent on exploration compared to their control group. Thus, FLX-males did not differ from CTR-males throughout the different time points during the observation period. FLX-females other the other hand, scored lower than CTR-females during the whole hour, but most prominently in the first 10- (*t* = 3.03, *p* = .009, *d* = 1.52), 20- (*t* = 4.38, *p* = < .001, *d* = 2.19) and 30- minutes (*U* = 12.00, *z* = − 2.10, *p* = .038, *r* = −.53). Similar results were revealed for the frequency of non-social exploration (females: *F*(5,70) = 0.23, *p* = .948, η_p_^2^ = .02; males: *F*(5,70) = 1.76, *p* = .132, η_p_^2^ = .11).

## 4. Discussion

In our study, we investigated how perinatal fluoxetine exposure affects adult social and non-social investigation behaviors in a novel seminatural environment with unfamiliar conspecifics. Our findings show that perinatal fluoxetine exposure does not induce alterations on social investigation behaviors and strategies when introduced to a novel seminatural environment and unknown conspecifics. However, perinatal fluoxetine exposure was found to affect non-social investigation behaviors. More specifically, perinatal fluoxetine exposed female and male rats showed increased locomotor activity (in terms of walking/running), while perinatal fluoxetine exposed females showed decreased non-social exploration. Furthermore, it was demonstrated that the observed differences maintained throughout the whole observation.

### 4.1 Social behaviors

The first question we investigated was whether social investigation behaviors, operationalized as active social behaviors (sniffing and grooming other rats) and latency to meet all other cohort members, would be affected by perinatal SSRI exposure. The ability to interact in line with social norms is crucial in everyday life, and deviant social behavior in the initial phase of contact can make it difficult to establish social relationships. The results in this study revealed no differences between CTR- and FLX-rats on the total time spent on, or the number of, active social behaviors. Previous findings from the same experiment showed that FLX-females, but not FLX-males, showed a tendency toward decreased active social behaviors (Houwing et al. 2019a), which was not present after naturally occurring aggressive encounters (Heinla et al. 2020). Nevertheless, in those studies, behaviors were observed after the rats had already been housed together in the seminatural environment for several days, and thus were familiar with each other. The effect of fluoxetine on social behaviors might have different outcomes depending on whether the rats are interacting with familiar or unfamiliar partners (Gemmel et al. 2019). In the present study, the rats were observed during the first hour after introduction to the seminatural environment, allowing us to investigate how the rats encounter the first social situations before knowing each other. It should be noted, however, that the combination of a novel environment together with novel conspecifics, might give different results than exposure to only one of the novelties. The use of a novel environment may have masked differences in active social behaviors, since such differences have been found in previous studies where animals were observed in a familiar environment (Heinla et al. 2020; Houwing et al. 2019a).

We also measured how long it took the rats to meet the other cohort members after being introduced to the novel environment. Such latency times could indicate whether the rats have different interests in approaching other rats. Lack of social interest is a relevant trait to examine since such symptoms commonly appear in various mental and neurodevelopmental disorders (Barkus and Badcock 2019). However, the results did not reveal any differences in latencies to meet conspecifics between CTR- and FLX-rats. From our findings, we conclude that perinatal SSRI exposure does not affect social investigation behavior and strategies during the first hour after introduction to a novel environment with unfamiliar conspecifics.

We further investigated whether SSRI exposure leads to behavioral alterations in other aspects of social behaviors, such as social passive behaviors and conflict behaviors. The results revealed no difference in passive social behavior between FLX-rats and CTR-rats. Furthermore, neither FLX-females nor FLX-males differed from CTR-rats in terms of conflict behavior. However, conflict behavior was not frequently occurring in our experiment. The Wistar strain is generally known to exhibit little aggressive behavior compared to other strains (Koolhaas et al. 2013). In addition, the experiment was not designed to trigger aggressive behavior as competition for food, water or mating partners were not necessary.

### 4.2 Non-social behaviors

Next, we investigated whether perinatal SSRI exposure would affect non-social investigation behaviors, operationalized as locomotor activity (walking/running), non-social exploration and latency to visit all locations of the environment. We found that both FLX-females and FLX-males spent more time on locomotor activity compared to control rats. The display of subtle changes in motor development and in motor movement has also been found in children exposed to SSRIs in (Casper et al. 2003). However, FLX-female rats also visited more locations of the seminatural environment within the first 3 minutes after entrance compared to CTR-females. This could indicate that perinatal SSRI exposure leads to an increased interest to investigate paths and locations. Contrary to our findings, a recent meta-analysis found evidence for *reduced* activity in developmentally SSRI exposed rats, as mostly measured by total distance moved (Ramsteijn et al. 2020). Although we did not measure total distance per se, it is reasonable to assume that total distance is related to total time spent walking/running in the seminatural environment. Nevertheless, the meta-analysis is mainly based on studies measuring activity in simplified open field boxes. Such set-ups allow the rats to perceive the whole environment without necessarily having to move their bodies. We could therefore assume that an increased interest to investigate locations and paths would only be observable in situations where walking/running (movement) is needed to investigate the environment. In addition, in the current environment, more rats were present leading to the assumption that the odors and sounds from others may also elicit extra movement, making our set-up more reliable to study the effects of perinatal fluoxetine exposure on a measure as locomotor activity reflecting alterations in interest to investigate novel paths and locations. With this in mind, we would expect the differences between FLX- and CTR-rats to disappear (or diminish) when the animals get familiar with their surroundings and conspecifics. Interestingly, previous studies from our research group that analyzed other relevant data from the same experiment did indeed find no differences on locomotor activity between FLX- and CTR-rats after the rats were already familiarized with the environment (Heinla et al. 2020; Houwing et al. 2019a). This suggests that the current findings of increased locomotor activity in FLX-rats are related to the introduction to a novel environment, and not the complexity of the environment on itself.

We also found that FLX-females, but not FLX-males, spent less time on non-social exploration than control rats, meaning they were sniffing less on objects (e.g. shelters, wooden sticks) and specific elements in the environment (e.g. walls, the ground). This is in line with previous findings from day 4 and day 7 in the same experiment (Houwing et al. 2019a), where reduced non-social exploration was found in FLX-females, but not in FLX-males. However, since in the present study males and females were not compared directly, any conclusion on the relative difference between the sexes can’t be drawn. Other studies have also reported reduced non-social explorative behaviors in SSRI exposed rats (Ansorge et al. 2004; Karpova et al. 2009; Rebello et al. 2014; Sarkar et al. 2014; Simpson et al. 2011; Zohar et al. 2016). Although we have shown that FLX-females seem to have increased interest to explore paths and locations, shown by increased locomotor activity, our findings also indicate that perinatal SSRI exposure in females leads to reduced interest to investigate objects and other specific elements in the environment. Although the findings might seem contradictive at first sight, locomotor activity and non-social exploration could possibly serve different purposes. As locomotor activity could measure the interest to get quickly familiar with the whole environment as a kind of screening behavior, non-social exploration reflects a more detailed and accurate investigation of the environment. Therefore, we suggest that perinatal SSRI exposure alters the strategy the animals use to investigate a novel environment leading to a quicker, but less detailed investigation of novel environments. Previous studies have found an association between perinatal SSRI exposure and diagnosis of attention-deficit hyperactivity disorder (ADHD; Man et al, 2018). It could be speculated that the investigation strategy observed in FLX-rats could reflect ADHD-related symptoms, such as often failing to give close attention to details and disliking tasks requiring sustained mental effort (American Psychiatric Association, 2013).

We further investigated other non-social behaviors such as anxiety/stress-related behaviors. We did not find any difference between CTR- and FLX-rats on anxiety/stress-related behaviors. Another subset of data from this experiment using the same cohorts of animals found that white-noise exposure induced increased self-grooming in FLX-males (Houwing et al. 2019a), which was explained as an altered stress-coping behavior. As introduction to a new environment can be considered a stressful situation, we expected to observe a similar increase in self-grooming behavior in FLX-males in the present study. However, no differences were found between CTR- and FLX-rats on self-grooming behavior. Moreover, no differences were found on the amount of time spent in the open area, as measure for changes in anxiety-related behavior. Altogether, this makes us to conclude that perinatal SSRI exposure does not affect anxiety/stress-related behavior during the first hour of exposure to a novel environment with unfamiliar conspecifics.

### 4.3 Behavioral adaption over time

The last question we investigated was whether perinatal SSRI exposed rats adapt differently to unfamiliarity (both environmental and socially) than their non-exposed conspecifics. Therefore, we split the observational data into six 10-minute time-bins in order to look at behavioral changes over time. As part of the familiarization process to a new environment, we generally expected to see adjustments in behavior during the first hour, such as decrease in general activity (Wilkinson et al. 2006). However, our main subject of interest was whether perinatal SSRI exposed rats adjusted their behavior in a different manner than controls.

Our results revealed that SSRI exposed animals adapted similarly to the novel environment as control animals. As discussed, FLX-females spent less time exploring objects and the physical environment, whereas both FLX-males and FLX-females spent more time on locomotor activity compared to CTR-rats. Those differences remained relatively stable throughout the first hour, meaning that FLX- and CTR-rats behaved differently, but adapted similarly to the novel environment over time (the increased locomotor activity remained higher during the full course of the observed hour). We conclude that perinatal fluoxetine exposed rats do not adapt their behaviors differently than controls during the first hour after introduction to the novel environment, instead the changes in non-social investigation behavior remain stable over time.

## 5. Conclusion

In summary, our data showed that perinatal SSRI exposure alters aspects of non-social investigation behaviors when introduced to a novel environment with unfamiliar conspecifics, but did not alter social investigation behaviors. Both FLX-males and FLX-females showed a higher amount of locomotor activity, while FLX-females visited more locations within the first three minutes, and spent less time exploring objects and specific elements in the physical environment. Perinatal fluoxetine exposure did not affect social behavior or how the animals adapted to the unfamiliar seminatural environment over time. Altogether, we conclude that perinatal SSRI exposure alters non-social investigation, to a quicker and less detailed strategy, when exposed to a novel environment, and that the alteration is most pronounced in females.

## Acknowledgements

Financial support was received from Helse Nord #PFP1295-16, Norway. We also would like to thank Ragnhild Osnes, Carina Sørensen, Nina Løvhaug, Katrine Harjo, and Remi Osnes for their excellent care of the animals.

