## Supplemental materials for "Effects of Perinatal Fluoxetine Exposure on Novelty-induced Social and Non-Social Investigation Behaviors in a Seminatural Environment"

[3] Regional Health Authority of North Norway

[*] These authors contributed equally.

**Table S1**

*Treatment Dams and Cage Distribution Offspring*

| Dams | Treatment | Number offspring | Male-Cage 1 | Male-Cage 2 | Female-Cage 3 | Female-Cage 4 | Female-Cage 5 | Female-Cage 6 |
|---|---|---|---|---|---|---|---|---|
| F1 | FLX | 7 | OM1<br>OM2<br>OM3 | | OF2<br>OF3 | OF1<br>OF4 | | |
| F2 | FLX | 9 | OM5<br>OM6<br>OM7 | OM4<br>OM8 | OF5<br>OF8 | OF6<br>OF7 | | |
| F3 | CTR | 13 | OM9<br>OM12 | OM10<br>OM11 | OF12<br>OF16 | OF9<br>OF15 | OF11<br>OF14 | OF10<br>OF13<br>OF17 |
| F4 | CTR | 6 | OM13<br>OM14<br>OM15 | | OF18<br>OF19<br>OF20 | | | |
| F5 | FLX | None | | | | | | |
| F6 | FLX | None | | | | | | |
| F7 | CTR | None | | | | | | |
| F8 | CTR | 15 | OM16<br>OM17 | OM20<br>OM21<br>OM25 | OF21<br>OF23 | OF22<br>OF24<br>OF25 | | |
| F9 | FLX | 8 | OM29<br>OM30<br>OM31 | | OF27<br>OF29 | OF26<br>OF28<br>OF30 | | |
| F10 | FLX | dead | | | | | | |

*Note.* M = male, F= female, CTR = methylcellulose, FLX = fluoxetine, OM = male offspring, OF = female offspring

**Figure S1**

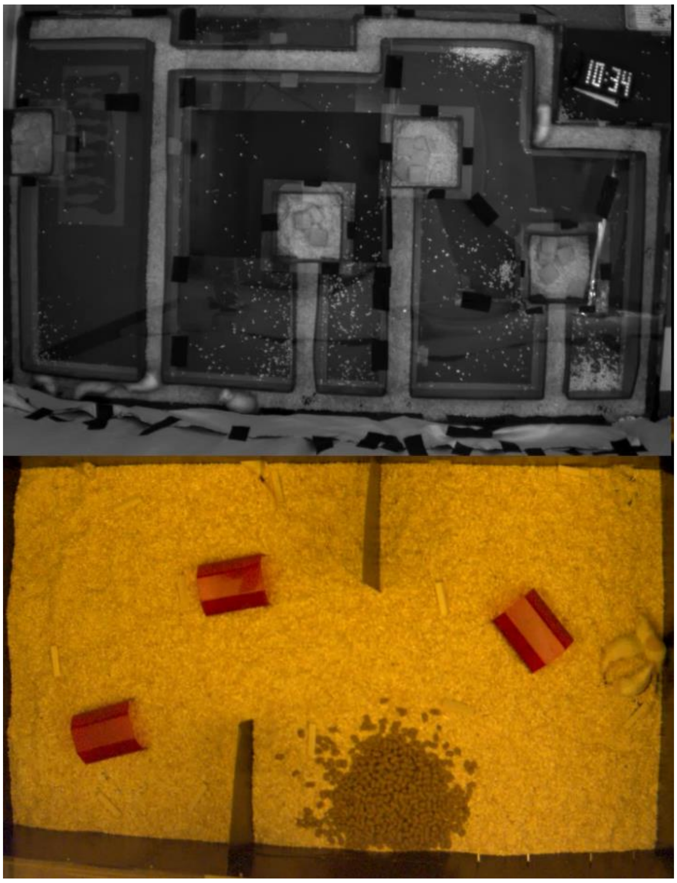

*Figure S1. Picture of the whole seminatural environment*

#### Table S2

*Experimental Design*

| Colony | Male 1 | Male 2 | Male 3 | Male 4 | Female 1 | Female 2 | Female 3 | Female 4 |
|---|---|---|---|---|---|---|---|---|
| SNE1 | OM1 | OM5 | OM9 | OM13 | OF2 | OF5 | OF12 | OF21 |
| SNE2 | OM2 | OM6 | OM12 | OM16 | OF3 | OF8 | OF13 | OF22 |
| SNE3 | OM3 | OM7 | OM15 | OM17 | OF1 | OF6 | OF15 | OF24 |
| SNE4 | OM4 | OM11 | OM21 | OM29 | OF4 | OF9 | OF19 | OF27 |
| SNE5 | OM8 | OM10 | OM20 | OM30 | OF7 | OF14 | OF20 | OF26 |

*Note.* SNE = seminatural environment, OM = male offspring, OF female offspring

#### Table S3

*Means and Standard Error for Duration of Behaviors*

| Behavior | | 10 | 20 | 30 | 40 | 50 | 60 | Total |
|---|---|---|---|---|---|---|---|---|
| Socially active behaviors | CTR-females | 80.5 ± 10.9 | 58.9 ± 6.1 | 61.4 ± 8.5 | 75.2 ± 9.5 | 69.2 ± 8.1 | 53.4 ± 5.8 | 398.6 ± 21.3 |
| | FLX-females | 89.0 ± 9.3 | 67.9 ± 9.0 | 65.9 ± 5.4 | 81.4 ± 9.5 | 86.4 ± 9.0 | 54.4 ± 10.5 | 445.1 ± 39.2 |
| | CTR-males | 97.7 ± 13.6 | 104.3 ± 23.6 | 120.6 ± 16.8 | 109.5 ± 17.2 | 110.3 ± 13.0 | 94.4 ± 16.9 | 636.5 ± 62.1 |
| | FLX-males | 79.7 ± 12.9 | 100.5 ± 25.3 | 91.2 ± 13.8 | 77.3 ± 10.9 | 106.0 ± 14.2 | 93.6 ± 13.0 | 548.0 ± 68.8 |
| General activity | CTR-females | 412.7 ± 11.0 | **400.2 ± 7.4** | 391.5 ± 11.6 | 391.4 ± 12.4 | 361.9 ± 12.7 | 388.3 ± 19.3 | 2346.0 ± 35.1 |
| | FLX-females | 391.5 ± 8.4 | **372.7 ± 8.6** | 394.9 ± 11.8 | 389.3 ± 5.7 | 368.3 ± 27.1 | 390.0 ± 8.0 | 2308.1 ± 51.2 |
| | CTR-males | 362.1 ± 28.6 | 339.4 ± 27.3 | 298.1 ± 18.0 | **267.7 ± 13.9** | 287.4 ± 28.9 | **256.4 ± 19.4** | 1810.3 ± 95.4 |
| | FLX-males | 383.1 ± 19.6 | 336.6 ± 24.0 | 325 ± 21.6 | **327.5 ± 9.5** | 323.9 ± 15.3 | **308.3 ± 14.3** | 2006.0 ± 66.1 |
| Non-socially passive behaviors | CTR-females | 16.4 ± 4.7 | 12.3 ± 3.3 | 14.3 ± 4.0 | 16.4 ± 4.4 | 36.8 ± 13.2 | 30.0 ± 10.0 | 568.1 ± 59.7 |
| | FLX-females | 12.7 ± 2.8 | 11.9 ± 2.1 | 15.6 ± 2.3 | 14.2 ± 5.0 | 25.3 ± 13.0 | 26.8 ± 4.5 | 614.7 ± 80.5 |
| | CTR-males | 21.1 ± 5.9 | 19.1 ± 5.1 | 27.9 ± 6.3 | 46.0 ± 14.1 | 39.4 ± 18.8 | 60.8 ± 16.8 | 273.7 ± 44.1 |
| | FLX-males | 18.5 ± 6.8 | 13.4 ± 3.1 | 17.8 ± 3.9 | 36.0 ± 11.6 | 14.5 ± 4.1 | 34.6 ± 10.2 | 163.0 ± 30.6 |
| Socially passive behaviors | CTR-females | 2.5 ± 1.4 | 6.3 ± 3.2 | 3.2 ± 1.1 | 4.2 ± 1.1 | 16.9 ± 8.5 | 5.1 ± 2.7 | 38.1 ± 9.3 |
| | FLX-females | 2.8 ± 1.5 | 3.9 ± 1.9 | 8.1 ± 3.4 | 4.9 ± 1.7 | 18.1 ± 10.6 | 3.8 ± 1.6 | 41.6 ± 14.2 |
| | CTR-males | 4.1 ± 2.2 | 3.3 ± 2.7 | 2.6 ± 1.6 | 15.4 ± 5.3 | 14.9 ± 8.2 | 19.3 ± 8.7 | 29.2 ± 10.0 |
| | FLX-males | 2.6 ± 1.3 | 2.5 ± 1.1 | 3.9 ± 1.3 | 7.3 ± 3.2 | 4.6 ± 1.4 | 7.7 ± 1.8 | 29.6 ± 6.7 |
| Conflict behaviors | CTR-females | 8.2 ± 2.6 | 5.9 ± 1.7 | 11.4 ± 4.1 | 14.1 ± 4.8 | 11.2 ± 3.3 | 8.1 ± 1.8 | 58.8 ± 12.0 |
| | FLX-females | 8.4 ± 2.0 | 12.6 ± 3.1 | 10.7 ± 3.0 | 7.9 ± 1.5 | 8.1 ± 2.5 | 10.9 ± 3.7 | 58.4 ± 9.8 |
| | CTR-males | 1.5 ± 0.9 | 1.9 ± 0.9 | 3.8 ± 1.7 | 9.3 ± 3.7 | 4.6 ± 2.6 | 19.5 ± 13.4 | 40.6 ± 16.5 |
| | FLX-males | 2.0 ± 1.1 | 4.2 ± 3.1 | 10.9 ± 4.0 | 16.5 ± 7.2 | 9.5 ± 2.2 | 8.7 ± 4.6 | 51.2 ± 13.2 |
| Walking/running | CTR-females | **74.1 ± 8.1** | 79.7 ± 14.7 | **63.2 ± 6.2** | 68.1 ± 7.7 | 65.4 ± 9.7 | 54.4 ± 7.9 | **404.9 ± 29.4** |
| | FLX-females | **111.6 ± 10.9** | 116.3 ± 14.7 | **115.2 ± 18.0** | 95.5 ± 10.5 | 102.2 ± 24.7 | 75.7 ± 13.1 | **616.5 ± 80.1** |
| | CTR-males | 157.3 ± 17.3 | 155.5 ± 13.1 | 131.8 ± 11.9 | **143.4 ± 6.4** | **129.0 ± 14.6** | 122.3 ± 6.7 | **839.0 ± 40.7** |
| | FLX-males | 176.7 ± 18.7 | 175.8 ± 14.0 | 162.8 ± 13.1 | **186.4 ± 10.2** | **185.5 ± 16.6** | 154.8 ± 13.7 | **1041.6 ± 52.6** |
| Chasing | CTR-females | 0.0 ± 0.0 | 0.0 ± 0.0 | 0.0 ± 0.0 | 0.3 ± 0.03 | 0.0 ± 0.0 | 0.0 ± 0.0 | 0.3 ± 0.3 |
| | FLX-females | 0.0 ± 0.0 | 0.0 ± 0.0 | 0.0 ± 0.0 | 0.0 ± 0.0 | 0.0 ± 0.0 | 0.0 ± 0.0 | 0.0 ± 0.0 |
| | CTR-males | 2.8 ± 1.8 | 0.2 ± 0.2 | 0.0 ± 0.0 | 0.0 ± 0.0 | 0.0 ± 0.0 | 0.0 ± 0.0 | 3.0 ± 2.0 |
| | FLX-males | 0.0 ± 0.0 | 0.3 ± 0.3 | 0.3 ± 0.3 | 3.6 ± 2.4 | 0.0 ± 0.0 | 1.0 ± 0.7 | 5.3 ± 3.2 |
| Non-social exploration | CTR-females | **338.7 ± 16.2** | **320.5 ± 8.8** | **328.3 ± 10.5** | 323.3 ± 18.0 | 296.4 ± 11.8 | 333.9 ± 23.1 | **1941.1 ± 48.0** |
| | FLX-females | **279.9 ± 10.7** | **256.3 ± 11.7** | **279.6 ±14.6** | 293.8 ± 13.1 | 266.1 ± 26.1 | 314.3 ± 15.4 | **1691.5 ± 64.7** |
| | CTR-males | 204.8 ± 36.1 | 183.9 ± 29.2 | 166.3 ± 16.9 | 124.3 ± 11.0 | 158.5 ± 22.5 | 134.0 ± 19.9 | 971.3 ± 89.5 |
| | FLX-males | 206.4 ± 31.6 | 160.9 ± 13.5 | 162.5 ± 13.9 | 141.0 ± 16.9 | 138.3 ± 14.6 | 153.5 ± 16.2 | 964.4 ± 72.0 |
| Digging | CTR-females | 5.5 ± 3.2 | 31.0 ± 11.6 | 26.0 ± 4.8 | 17.8 ± 5.6 | 14.7 ± 4.1 | 16.9 ± 4.8 | 111.8 ± 21.9 |
| | FLX-females | 12.6 ± 3.5 | 28.0 ± 7.6 | 32.2 ± 7.5 | 14.6 ± 2.2 | 15.7 ± 2.2 | 29.9 ± 8.1 | 133.1 ± 18.3 |
| | CTR-males | 0.8 ± 0.8 | 2.4 ± 1.7 | 16.7 ± 8.3 | 8.5 ± 7.0 | 3.0 ± 1.8 | 2.9 ± 1.9 | 34.2 ± 14.4 |
| | FLX-males | 4.7 ± 2.8 | 6.4 ± 3.0 | 13.3 ± 8.7 | 5.3 ± 4.0 | 2.9 ± 1.8 | 3.6 ± 2.2 | 33.9 ± 14.6 |
| Resting/immobile alone | CTR-females | 1.1 ± 0.7 | 0.0 ± 0.0 | 3.6 ± 3.3 | 0.5 ± 0.3 | 7.9 ± 6.4 | 8.8 ± 8.3 | 21.8 ± 14.7 |
| | FLX-females | 1.0 ± 1.0 | 0.3 ± 0.3 | 3.1 ± 1.9 | 4.0 ± 2.6 | 4.6 ± 3.8 | 11.8 ± 4.4 | 24.8 ± 9.2 |
| | CTR-males | 17.5 ± 6.4 | 16.4 ± 5.6 | 24.7 ± 6.6 | 34.9 ± 8.6 | 33.5 ± 17.5 | 57.1 ± 17.5 | 183.8 ± 35.4 |
| | FLX-males | 15.3 ± 6.6 | 11.8 ± 3.1 | 14.6 ± 4.2 | 32.2 ± 11.7 | 13.4 ± 4.1 | 29.1 ± 10.1 | 116.1 ± 27.6 |
| Resting/immobile socially | CTR-females | 0.0 ± 0.0 | 0.0 ± 0.0 | 0.0 ± 0.0 | 0.3 ± 0.3 | 0.8 ± 0.4 | 0.0 ± 0.0 | 1.1 ± 0.6 |
| | FLX-females | 0.0 ± 0.0 | 0.0 ± 0.0 | 0.2 ± 0.2 | 1.2 ± 0.7 | 6.4 ± 4.0 | 0.2 ± 0.2 | 7.9 ± 4.5 |

| | | | | | | | | |
|---|---|---|---|---|---|---|---|---|
| | CTR-males | 2.0 ± 0.9 | 1.4 ± 0.8 | 0.4 ± 0.3 | 9.5 ± 4.9 | 14.1 ± 8.4 | 18.2 ± 9.0 | 45.5 ± 19.9 |
| | FLX-males | 2.4 ± 1.4 | 0.7 ± 0.5 | 2.4 ± 1.2 | 4.7 ± 2.9 | 2.8 ± 1.2 | 5.0 ± 1.5 | 18.1 ± 4.9 |
| Hiding alone | CTR-females | 15.3 ± 4.6 | 12.3 ± 3.3 | 10.7 ± 3.1 | 16.0 ± 4.5 | 28.8 ± 9.6 | 21.3 ± 5.7 | 104.3 ± 18.8 |
| | FLX-females | 11.7 ± 2.2 | 11.6 ± 2.2 | 12.5 ± 2.5 | 10.2 ± 3.9 | 20.7 ± 9.4 | 15.0 ± 4.1 | 81.7 ± 11.7 |
| | CTR-males | 3.5 ± 2.1 | 2.8 ± 1.1 | 3.2 ± 2.2 | 11.1 ± 10.1 | 5.9 ± 2.1 | 3.7 ± 2.2 | 30.2 ± 15.1 |
| | FLX-males | 3.2 ± 1.5 | 1.6 ± 0.9 | 3.2 ± 1.7 | 3.8 ± 1.4 | 1.1 ± 0.5 | 5.5 ± 3.2 | 18.4 ± 5.1 |
| Hiding socially | CTR-females | 2.5 ± 1.4 | 6.3 ± 3.2 | 3.2 ± 1.1 | 3.9 ± 1.2 | 16.1 ± 8.4 | 5.1 ± 2.7 | 37.1 ± 9.4 |
| | FLX-females | 2.8 ± 1.5 | 3.9 ± 1.9 | 7.9 ± 3.4 | 3.7 ± 1.9 | 11.8 ± 8.3 | 3.6 ± 1.6 | 33.7 ± 12.7 |
| | CTR-males | 2.1 ± 2.1 | 2.0 ± 2.0 | 2.3 ± 1.7 | 5.9 ± 2.5 | 0.8 ± 0.5 | 1.1 ± 0.8 | 14.1 ± 5.8 |
| | FLX-males | 1.9 ± 1.9 | 1.8 ± 1.2 | 1.5 ± 0.9 | 2.5 ± 1.5 | 1.8 ± 1.4 | 2.8 ± 1.2 | 10.5 ± 4.1 |
| Following | CTR-females | 8.2 ± 6.1 | 1.7 ± 1.3 | 1.7 ± 1.0 | 1.9 ± 1.4 | 1.3 ± 1.0 | 1.6 ± 1.3 | 16.5 ± 10.7 |
| | FLX-females | 4.6 ± 1.9 | 2.3 ± 1.4 | 1.6 ± 0.6 | 1.8 ± 1.2 | 1.5 ± 1.0 | 1.3 ± 0.9 | 13.0 ± 5.0 |
| | CTR-males | 33.3 ± 9.0 | 30.8 ± 7.8 | 36.5 ± 10.3 | 48.8 ± 12.7 | 39.7 ± 8.5 | 36.5 ± 10.5 | 225.5 ± 47.9 |
| | FLX-males | 30.7 ± 5.6 | 36.5 ± 6.7 | 34.4 ± 5.3 | 26.1 ± 6.0 | 32.2 ± 6.5 | 30.6 ± 5.9 | 189.7 ± 26.4 |
| Allogrooming | CTR-females | 0.0 ± 0.0 | 0.0 ± 0.0 | 0.0 ± 0.0 | 1.0 ± 0.8 | 1.4 ± 1.1 | 0.2 ± 0.2 | 2.6 ± 1.1 |
| | FLX-females | 0.0 ± 0.0 | 0.3 ± 0.3 | 0.0 ± 0.0 | 0.0 ± 0.0 | 0.9 ± 0.9 | 0.2 ± 0.2 | 1.3 ± 0.9 |
| | CTR-males | 1.5 ± 1.0 | 3.7 ± 3.0 | 1.0 ± 1.0 | 2.3 ± 1.6 | 1.8 ± 1.5 | 4.4 ± 4.4 | 14.5 ± 12.1 |
| | FLX-males | 1.2 ± 0.8 | 1.2 ± 1.2 | 0.0 ± 0.0 | 0.2 ± 0.2 | 0.7 ± 0.4 | 0.2 ± 0.2 | 3.5 ± 2.0 |
| Sniffing anogenitally | CTR-females | 35.9 ± 9.0 | 20.7 ± 2.3 | 20.1 ± 5.8 | 19.0 ± 5.4 | 20.9 ± 3.6 | 14.8 ± 2.2 | 131.4 ± 8.3 |
| | FLX-females | 39.7 ± 5.4 | 27.7 ± 5.8 | 23.5 ± 5.1 | 29.6 ± 7.7 | 25.2 ± 5.1 | 17.7 ± 3.7 | 163.1 ± 12.3 |
| | CTR-males | 52.9 ± 12.1 | 64.1 ± 19.7 | 85.9 ± 14.7 | 68.7 ± 13.2 | 68.1 ± 9.6 | 52.5 ± 11.6 | 391.9 ± 44.6 |
| | FLX-males | 38.2 ± 10.7 | 70.8 ± 21.1 | 62.3 ± 14.4 | 43.8 ± 10.4 | 66.3 ± 13.2 | 61.1 ± 8.9 | 342.3 ± 58.3 |
| Sniffing nose-to-nose | CTR-females | 13.5 ± 5.0 | 10.5 ± 3.8 | 16.3 ± 8.2 | 21.4 ± 4.2 | 18.1 ± 1.7 | 15.7 ± 2.6 | 95.5 ± 21.9 |
| | FLX-females | 10.4 ± 2.3 | 9.4 ± 0.8 | 14.5 ± 2.2 | 18.3 ± 3.1 | 27.5 ± 10.8 | 10.3 ± 2.0 | 90.6 ± 15.4 |
| | CTR-males | 3.5 ± 0.7 | 3.6 ± 1.1 | 4.6 ± 1.5 | 7.5 ± 2.2 | 8.8 ± 3.7 | 8.8 ± 1.8 | 36.7 ± 9.0 |
| | FLX-males | 3.6 ± 1.2 | 2.2 ± 1.1 | 3.5 ± 1.0 | 5.0 ± 0.9 | 6.4 ± 2.3 | 5.7 ± 1.3 | 26.4 ± 5.6 |
| All sniffing | CTR-females | 80.5 ± 10.9 | 58.9 ± 6.1 | 61.4 ± 8.5 | 74.3 ± 9.5 | 67.8 ± 7.4 | 53.1 ± 5.8 | 360.0 ± 22.2 |
| | FLX-females | 89.0 ± 9.5 | 67.6 ± 8.8 | 65.9 ± 5.4 | 81.4 ± 9.5 | 85.6 ± 9.2 | 54.3 ± 5.8 | 443.8 ± 39.2 |
| | CTR-males | 96.2 ± 14.4 | 100.6 ± 23.4 | 119.6 ± 16.5 | 107.2 ± 16.7 | 108.6 ± 12.4 | 90.0 ± 14.3 | 622.0 ± 54.8 |
| | FLX-males | 78.5 ± 13.1 | 99.3 ± 24.4 | 91.2 ± 13.8 | 77.1 ± 11.0 | 105.3 ± 14.4 | 93.4 ± 13.0 | 544.5 ± 68.8 |
| Sniffing body | CTR-females | 31.2 ± 5.3 | 27.7 ± 5.1 | 25.0 ± 3.9 | 33.8 ± 6.4 | 28.8 ± 5.0 | 22.6 ± 3.0 | 169.1 ± 21.6 |
| | FLX-females | 39.1 ± 6.0 | 30.5 ± 3.7 | 27.9 ± 3.9 | 33.5 ± 3.5 | 32.9 ± 5.4 | 26.2 ± 6.3 | 190.1 ± 21.7 |
| | CTR-males | 40.0 ± 3.0 | 33.0 ± 4.8 | 29.2 ± 6.5 | 31.0 ± 4.6 | 31.7 ± 6.7 | 28.7 ± 4.1 | 193.4 ± 12.5 |
| | FLX-males | 36.8 ± 4.9 | 26.3 ± 4.1 | 25.3 ± 2.7 | 28.2 ± 4.8 | 32.6 ± 3.0 | 26.7 ± 4.9 | 175.7 ± 18.6 |
| Fighting | CTR-females | 8.2 ± 2.6 | 5.9 ± 1.7 | 11.4 ± 4.1 | 14.1 ± 4.8 | 11.2 ± 3.3 | 8.1 ± 1.8 | 58.8 ± 12.0 |
| | FLX-females | 8.4 ± 2.0 | 12.6 ± 3.1 | 10.7 ± 3.0 | 7.8 ± 1.5 | 8.1 ± 2.5 | 10.9 ± 3.7 | 58.4 ± 9.8 |
| | CTR-males | 1.5 ± 0.9 | 1.9 ± 0.9 | 3.8 ± 1.7 | 8.3 ± 3.3 | 4.2 ± 2.5 | 5.8 ± 2.1 | 25.5 ± 6.0 |
| | FLX-males | 2.0 ± 1.1 | 4.2 ± 3.1 | 10.9 ± 4.0 | 15.1 ± 6.7 | 7.2 ± 2.0 | 5.8 ± 2.8 | 45.2 ± 11.9 |
| Nose-off | CTR-females | 0.0 ± 0.0 | 0.0 ± 0.0 | 0.0 ± 0.0 | 0.0 ± 0.0 | 0.0 ± 0.0 | 0.0 ± 0.0 | 0.0 ± 0.0 |
| | FLX-females | 0.0 ± 0.0 | 0.0 ± 0.0 | 0.0 ± 0.0 | 0.1 ± 0.1 | 0.0 ± 0.0 | 0.0 ± 0.0 | 0.1 ± 0.1 |
| | CTR-males | 0.0 ± 0.0 | 0.0 ± 0.0 | 0.0 ± 0.0 | 1.9 ± 0.6 | 0.4 ± 0.2 | 13.7 ± 12.4 | 15.1 ± 12.7 |
| | FLX-males | 0.0 ± 0.0 | 0.0 ± 0.0 | 0.1 ± 01 | 1.3 ± 0.6 | 2.3 ± 1.3 | 2.9 ± 1.9 | 6.0 ± 2.4 |
| Self-grooming | CTR-females | 8.5 ± 3.2 | 5.3 ± 1.6 | 20.2 ± 5.8 | 12.1 ± 3.4 | 19.0 ± 7.6 | 29.1 ± 10.9 | 94.2 ± 12.5 |
| | FLX-females | 3.4 ± 1.1 | 6.9 ± 1.3 | 8.8 ± 2.4 | 15.0 ± 5.1 | 11.2 ± 3.1 | 25.3 ± 7.4 | 70.5 ± 6.8 |
| | CTR-males | 1.0 ± 0.6 | 7.3 ± 4.4 | 3.4 ± 2.3 | 9.6 ± 5.0 | 21.9 ± 11.7 | 22.3 ± 9.5 | 65.2 ± 25.3 |
| | FLX-males | 1.2 ± 0.8 | 2.6 ± 1.1 | 16.7 ± 5.8 | 19.0 ± 9.5 | 10.3 ± 5.8 | 15.1 ± 6.0 | 64.6 ± 16.5 |
| Freezing | CTR-females | 0.0 ± 0.0 | 0.1 ± 0.1 | 0.4 ± 0.2 | 5.3 ± 3.2 | 1.6 ± 0.8 | 0.4 ±0.2 | 7.7 ± 3.6 |
| | FLX-females | 0.0 ± 0.0 | 0.1 ± 0.1 | 0.6 ± 0.4 | 6.9 ± 4.1 | 2.9 ± 1.1 | 0.9 ± 0.4 | 11.5 ± 4.6 |
| | CTR-males | 0.9 ± 0.6 | 0.0 ± 0.0 | 0.2 ± 0.2 | 3.6 ± 1.9 | 0.5 ± 0.3 | 0.9 ± 0.5 | 6.0 ± 2.3 |
| | FLX-males | 0.2 ± 0.2 | 0.0 ± 0.0 | 0.6 ± 0.6 | 4.0 ± 2.6 | 0.3 ± 0.2 | 7.1 ± 3.0 | 12.1 ± 3.4 |
| Rearing supported | CTR-females | 41.7 ± 5.9 | 62.5 ± 6.2 | 52.3 ± 8.7 | 47.7 ± 9.3 | 53.8 ± 7.6 | 58.2 ± 8.8 | 316.2 ± 25.3 |
| | FLX-females | 60.3 ± 8.7 | 80.8 ± 9.2 | 51.3 ± 5.2 | 51.4 ± 10.4 | 50.1 ± 5.9 | 49.1 ± 6.3 | 343.2 ± 27.6 |
| | CTR-males | 20.9 ± 3.0 | 37.7 ± 8.1 | 26.8 ± 4.2 | 24.8 ± 6.6 | 16.7 ± 4.9 | 19.5 ± 4.3 | 145.9 ± 17.5 |
| | FLX-males | 20.9 ± 3.4 | 36.4 ± 7.5 | 26.1 ± 6.1 | 16.9 ± 5.4 | 16.0 ± 4.7 | 13.5 ± 2.1 | 129.8 ± 18.4 |
| Rearing unsupported | CTR-females | 0.9 ± 0.2 | 1.9 ± 1.2 | 1.8 ± 0.7 | 1.5 ± 0.5 | 3.6 ± 1.2 | 1.9 ± 0.6 | 11.4 ± 3.1 |
| | FLX-females | 0.7 ± 0.3 | 2.5 ± 1.4 | 2.1 ± 1.9 | 2.1 ± 0.9 | 3.9 ± 1.2 | 3.0 ± 1.2 | 14.3 ± 2.6 |
| | CTR-males | 0.5 ± 0.4 | 0.5 ± 0.3 | 3.9 ± 1.8 | 4.6 ± 1.9 | 1.7 ± 0.8 | 3.8 ± 1.8 | 15.0 ± 5.6 |
| | FLX-males | 0.3 ± 0.3 | 1.7 ± 1.3 | 4.4 ± 1.8 | 1.6 ± 0.7 | 1.6 ± 0.7 | 9.7 ± 4.0 | 19.1 ± 3.6 |

*Note.* The data represent the time spent (s) performing all behaviors measured within the six time-bins. Data are shown in mean ± standard error of the mean.

### Table S4

*Means and Standard Error for Frequency of Behaviors*

| Behavior | | 10 | 20 | 30 | 40 | 50 | 60 | Total |
|---|---|---|---|---|---|---|---|---|
| Socially active behaviors | CTR-females | 59.0 ± 10.7 | 55.1 ± 7.0 | 49.4 ± 4.9 | 66.9 ± 9.3 | 56.5 ± 5.3 | 48.5 ± 3.6 | 335.4 ± 32.0 |
| | FLX-females | 69.5 ± 6.6 | 68.1 ± 8.7 | 62.8 ± 5.1 | 67.4 ± 5.8 | 66.5 ± 9.2 | 48.3 ± 8.2 | 382.6 ± 32.5 |
| | CTR-males | 47.9 ± 8.7 | 44.6 ± 7.0 | 54.0 ± 8.5 | 51.1 ± 8.1 | 50.5 ± 5.1 | 51.0 ± 9.2 | 299.0 ± 36.5 |
| | FLX-males | 42.4 ± 6.7 | 45.1 ± 8.7 | 46.9 ± 3.9 | 47.0 ± 5.3 | 61.4 ± 6.6 | 51.3 ± 4.9 | 294.0 ± 31.6 |
| General activity | CTR-females | 159.0 ± 24.1 | 175.5 ± 22.1 | 171.4 ± 21.9 | 177.4 ± 18.5 | 162.6 ± 17.6 | 155.9 ± 9.3 | 1001.8 ± 93.2 |
| | FLX-females | 204.8 ± 17.8 | 224.3 ± 14.9 | 213.8 ± 9.3 | 207.4 ± 15.4 | 189.8 ± 16.9 | 169.3 ± 22.8 | 1209.6 ± 66.1 |
| | CTR-males | 92.5 ± 6.4 | 98.6 ± 9.6 | 88.8 ± 10.0 | 83.9 ± 8.8 | 76.8 ± 8.3 | 75.5 ± 9.4 | 516.0 ± 44.2 |
| | FLX-males | 97.3 ± 6.9 | 99.3 ± 4.9 | 100.5 ± 5.3 | 98.3 ± 5.2 | 97.6 ± 6.1 | 97.5 ± 5.3 | 590.3 ± 13.6 |
| Non-socially passive behaviors | CTR-females | 5.5 ± 0.3 | 5.6 ± 1.0 | 5.6 ± 1.6 | 6.6 ± 1.5 | 7.8 ± 2.0 | 8.0 ± 2.3 | 51.6 ± 8.1 |
| | FLX-females | 5.5 ± 1.0 | 5.5 ± 0.9 | 7.4 ± 1.3 | 6.4 ± 1.5 | 7.3 ± 1.6 | 8.0 ± 1.7 | 55.0 ± 6.2 |
| | CTR-males | 3.9 ± 0.8 | 3.9 ± 0.6 | 4.1 ± 1.1 | 6.3 ± 1.6 | 7.0 ± 2.6 | 8.0 ± 1.5 | 44.6 ± 5.0 |
| | FLX-males | 4.3 ± 1.6 | 2.9 ± 0.6 | 4.0 ± 0.7 | 6.5 ± 1.7 | 3.9 ± 1.0 | 7.3 ± 2.0 | 35.5 ± 5.9 |
| Socially passive behaviors | CTR-females | 1.0 ± 0.3 | 2.0 ± 0.6 | 1.4 ± 0.5 | 1.9 ± 0.4 | 3.8 ± 1.2 | 2.0 ± 0.9 | 12.0 ± 2.7 |
| | FLX-females | 1.1 ± 0.4 | 1.8 ± 0.7 | 2.5 ± 0.8 | 1.5 ± 0.4 | 4.5 ± 1.4 | 1.6 ± 0.7 | 13.0 ± 2.4 |
| | CTR-males | 1.3 ± 0.8 | 0.9 ± 0.6 | 0.5 ± 0.2 | 2.4 ± 0.7 | 3.6 ± 1.6 | 2.9 ± 1.1 | 11.5 ± 3.1 |
| | FLX-males | 0.5 ± 1.2 | 0.6 ± 0.3 | 1.0 ± 0.3 | 1.6 ± 0.7 | 1.3 ± 0.3 | 1.8 ± 0.3 | 6.8 ± 0.9 |
| Conflict behaviors | CTR-females | 3.8 ± 0.9 | 5.0 ± 1.2 | 4.9 ± 1.2 | 7.8 ± 1.0 | 5.8 ± 1.9 | 5.0 ± 0.8 | 32.1 ± 3.9 |
| | FLX-females | 5.4 ± 1.5 | 6.5 ± 1.6 | 6.5 ± 1.3 | 6.5 ± 1.1 | 4.9 ± 1.1 | 5.0 ± 1.8 | 34.8 ± 5.2 |
| | CTR-males | 0.8 ± 0.4 | 1.4 ± 0.5 | 1.6 ± 0.5 | 3.1 ± 1.1 | 2.6 ± 0.9 | 1.9 ± 0.8 | 11.4 ± 2.7 |
| | FLX-males | 0.9 ± 0.5 | 1.1 ± 0.5 | 2.8 ± 0.5 | 6.3 ± 2.7 | 3.9 ± 1.1 | 2.4 ± 0.9 | 17.1 ± 4.0 |
| Walking/running | CTR-females | **44.4 ± 7.6** | **52.6 ± 8.5** | **51.1 ± 8.1** | **50.4 ± 6.5** | 46.1 ± 7.2 | 46.0 ± 4.5 | **290.6 ± 28.3** |
| | FLX-females | **75.4 ± 3.0** | **88.5 ± 8.9** | **81.6 ± 4.7** | **73.3 ± 5.3** | 71.6 ± 9.9 | 56.6 ± 10.7 | **447.0 ± 23.0** |
| | CTR-males | 52.5 ± 5.4 | 57.8 ± 6.9 | 51.8 ± 7.3 | 53.3 ± 6.2 | 46.6 ± 5.1 | **48.9 ± 6.9** | 310.8 ± 29.3 |
| | FLX-males | 54.6 ± 6.4 | 61.6 ± 4.3 | 61.5 ± 4.0 | 65.4 ± 5.1 | 64.4 ± 5.8 | **59.1 ± 4.5** | 366.4 ± 18.5 |
| Chasing | CTR-females | 0.0 ± 0.0 | 0.0 ± 0.0 | 0.0 ± 0.0 | 0.1 ± 0.1 | 0.0 ± 0.0 | 0.0 ± 0.0 | 0.1 ± 0.1 |
| | FLX-females | 0.0 ± 0.0 | 0.0 ± 0.0 | 0.0 ± 0.0 | 0.0 ± 0.0 | 0.0 ± 0.0 | 0.0 ± 0.0 | 0.0 ± 0.0 |
| | CTR-males | 1.8 ± 1.3 | 0.1 ± 0.1 | 0.0 ± 0.0 | 0.0 ± 0.0 | 0.0 ± 0.0 | 0.0 ± 0.0 | 1.9 ± 1.4 |
| | FLX-males | 0.0 ± 0.0 | 0.1 ± 0.1 | 0.1 ± 0.1 | 1.4 ± 0.9 | 0.0 ± 0.0 | 0.5 ± 0.4 | 2.1 ± 1.3 |
| Non-social exploration | CTR-females | 114.6 ± 17.0 | 122.9 ± 15.8 | 120.3 ± 14.4 | 127.0 ± 13.9 | 116.5 ± 11.2 | 109.9 ± 8.9 | 711.1 ± 71.1 |
| | FLX-females | 129.4 ± 15.0 | 135.8 ± 12.3 | 132.1 ± 12.0 | 134.1 ± 12.9 | 118.1 ± 13.9 | 112.6 ± 13.7 | 762.6 ± 65.0 |
| | CTR-males | 40.0 ± 3.8 | 40.9 ± 5.8 | 37.0 ± 3.8 | 30.6 ± 4.1 | 30.1 ± 4.9 | **26.6 ± 3.4** | 205.3 ± 21.7 |
| | FLX-males | 42.6 ± 3.3 | 37.6 ± 2.8 | 39.0 ± 2.7 | 32.9 ± 2.9 | 33.3 ± 3.3 | **38.4 ± 4.2** | 223.9 ± 13.7 |
| Digging | CTR-females | 2.9 ± 1.4 | 6.4 ± 1.9 | 6.1 ± 1.5 | 3.5 ± 1.1 | 3.4 ± 0.7 | 5.4 ± 1.1 | 27.6 ± 5.2 |
| | FLX-females | 5.0 ± 1.7 | 8.6 ± 1.7 | 8.8 ± 2.2 | 2.5 ± 0.7 | 3.4 ± 0.5 | 6.5 ± 1.3 | 35.8 ± 4.8 |
| | CTR-males | 0.3 ± 0.3 | 0.6 ± 0.4 | 0.9 ± 0.4 | 1.4 ± 1.0 | 0.9 ± 0.5 | 0.8 ± 0.5 | 4.8 ± 2.1 |
| | FLX-males | 0.9 ± 0.5 | 1.6 ± 0.7 | 2.5 ± 1.4 | 1.4 ± 0.8 | 0.5 ± 0.3 | 0.4 ± 0.2 | 7.1 ± 2.8 |
| Resting/immobile alone | CTR-females | 0.4 ± 0.3 | 0.0 ± 0.0 | 0.4 ± 0.3 | 0.5 ± 0.3 | 0.9 ± 0.4 | 0.4 ± 0.3 | 2.5 ± 1.0 |
| | FLX-females | 0.3 ± 0.3 | 0.1 ± 0.1 | 1.0 ± 0.6 | 1.5 ± 0.9 | 1.1 ± 0.5 | 2.1 ± 0.9 | 6.1 ± 2.2 |
| | CTR-males | 3.1 ± 0.9 | 3.4 ± 0.6 | 3.6 ± 1.1 | 5.3 ± 1.1 | 5.6 ± 2.2 | 7.4 ± 1.6 | 28.4 ± 4.2 |
| | FLX-males | 3.4 ± 1.5 | 2.3 ± 0.6 | 3.3 ± 0.7 | 5.5 ± 1.6 | 3.4 ± 1.0 | 6.0 ± 1.9 | 23.8 ± 5.1 |
| Resting/immobile socially | CTR-females | 0.0 ± 0.0 | 0.0 ± 0.0 | 0.0 ± 0.0 | 0.1 ± 0.1 | 0.4 ± 0.2 | 0.0 ± 0.0 | 0.5 ± 0.3 |
| | FLX-females | 0.0 ± 0.0 | 0.0 ± 0.0 | 0.1 ± 0.1 | 0.5 ± 0.3 | 1.3 ± 0.7 | 0.1 ± 0.1 | 2.0 ± 0.8 |
| | CTR-males | 0.5 ± 0.2 | 0.5 ± 0.3 | 0.3 ± 0.2 | 1.4 ± 0.7 | 3.3 ± 1.7 | 2.5 ± 1.2 | 8.4 ± 2.9 |
| | FLX-males | 0.4 ± 0.2 | 0.3 ± 0.2 | 0.5 ± 0.2 | 1.0 ± 0.6 | 0.9 ± 0.4 | 1.1 ± 0.2 | 4.1 ± 1.0 |
| Hiding alone | CTR-females | 5.1 ± 0.5 | 5.6 ± 1.0 | 5.3 ± 1.6 | 6.1 ± 1.6 | 6.9 ± 1.9 | 7.6 ± 2.2 | 36.6 ± 5.9 |
| | FLX-females | 5.3 ± 1.0 | 5.4 ± 0.9 | 6.4 ± 1.3 | 4.9 ± 1.4 | 6.1 ± 1.2 | 5.9 ± 1.7 | 33.9 ± 4.1 |
| | CTR-males | 0.8 ± 0.4 | 0.5 ± 0.2 | 0.5 ± 0.3 | 1.0 ± 0.7 | 1.4 ± 0.5 | 0.6 ± 0.3 | 4.8 ± 1.5 |
| | FLX-males | 0.9 ± 0.4 | 0.6 ± 0.3 | 0.8 ± 0.4 | 1.0 ± 0.3 | 0.5 ± 0.2 | 1.3 ± 0.7 | 5.0 ± 1.4 |
| Hiding socially | CTR-females | 1.0 ± 0.3 | 2.0 ± 0.6 | 1.4 ± 0.5 | 1.8 ± 0.5 | 3.4 ± 1.1 | 2.0 ± 0.9 | 11.5 ± 2.8 |
| | FLX-females | 1.1 ± 0.4 | 1.8 ± 0.7 | 2.4 ± 0.8 | 1.0 ± 0.4 | 3.3 ± 1.2 | 1.5 ± 0.7 | 11.0 ± 2.4 |
| | CTR-males | 0.8 ± 0.8 | 0.4 ± 0.4 | 0.3 ± 0.2 | 1.0 ± 0.3 | 0.4 ± 0.3 | 0.4 ± 0.3 | 3.1 ± 1.5 |
| | FLX-males | 0.1 ± 0.1 | 0.4 ± 0.3 | 0.5 ± 0.3 | 0.6 ± 0.3 | 0.4 ± 0.3 | 0.6 ± 0.3 | 2.6 ± 1.0 |
| Following | CTR-females | 2.0 ± 0.8 | 0.8 ± 0.5 | 0.6 ± 0.3 | 0.9 ± 0.4 | 0.6 ± 0.4 | 0.8 ± 0.5 | 5.6 ± 2.1 |
| | FLX-females | 1.9 ± 0.7 | 1.3 ± 0.6 | 1.1 ± 0.6 | 0.8 ± 0.5 | 1.1 ± 0.7 | 0.9 ± 0.5 | 7.0 ± 2.5 |
| | CTR-males | 16.6 ± 4.9 | 14.0 ± 3.2 | 17.9 ± 4.9 | 20.1 ± 5.3 | 16.9 ± 4.0 | 15.3 ± 5.1 | 100.8 ± 22.7 |
| | FLX-males | 12.9 ± 2.9 | 15.1 ± 3.1 | 14.0 ± 1.8 | 12.0 ± 2.5 | 13.4 ± 2.4 | 13.6 ± 2.8 | 81.1 ± 11.3 |

| | | | | | | | | |
|---|---|---|---|---|---|---|---|---|
| Allogrooming | CTR-females | 0.0 ± 0.0 | 0.0 ± 0.0 | 0.0 ± 0.0 | 0.4 ± 0.3 | 0.3 ± 0.2 | 0.1 ± 0.1 | 0.8 ± 0.3 |
| | FLX-females | 0.0 ± 0.0 | 0.1 ± 0.1 | 0.0 ± 0.0 | 0.0 ± 0.0 | 0.1 ± 0.1 | 0.1 ± 0.1 | 0.4 ± 0.2 |
| | CTR-males | 0.4 ± 0.3 | 0.6 ± 0.5 | 0.3 ± 0.3 | 0.3 ± 0.2 | 0.3 ± 0.2 | 0.8 ± 0.8 | 2.5 ± 2.0 |
| | FLX-males | 0.4 ± 0.3 | 0.4 ± 0.4 | 0.0 ± 0.0 | 0.1 ± 0.1 | 0.4 ± 0.2 | 0.1 ± 0.1 | 1.4 ± 0.8 |
| Sniffing anogenitally | CTR-females | 20.8 ± 5.1 | 15.6 ± 1.6 | 13.1 ± 3.1 | 13.6 ± 2.6 | 12.6 ± 1.6 | 9.9 ± 1.0 | 85.6 ± 7.0 |
| | FLX-females | 22.9 ± 2.5 | 20.3 ± 3.4 | 17.1 ± 2.8 | 16.6 ± 2.8 | 15.8 ± 3.0 | 11.8 ± 2.3 | 104.4 ± 7.0 |
| | CTR-males | 19.5 ± 5.2 | 20.1 ± 4.4 | 28.0 ± 5.8 | 23.3 ± 5.5 | 21.9 ± 3.9 | 20.4 ± 5.8 | 133.0 ± 22.8 |
| | FLX-males | 14.4 ± 3.4 | 21.5 ± 5.4 | 20.8 ± 3.4 | 16.3 ± 3.1 | 22.9 ± 3.4 | 20.5 ± 2.8 | 116.3 ± 17.7 |
| Sniffing nose-to-nose | CTR-females | 11.4 ± 1.9 | 11.5 ± 2.1 | 12.6 ± 1.8 | 21.1 ± 2.7 | 16.6 ± 1.1 | 16.9 ± 2.2 | 85.6 ± 7.0 |
| | FLX-females | 11.5 ± 1.1 | 12.1 ± 1.2 | 15.4 ± 1.4 | 19.8 ± 3.3 | 18.0 ± 1.7 | 12.1 ± 1.8 | 104.4 ± 7.0 |
| | CTR-males | 4.4 ± 0.8 | 4.8 ± 1.3 | 4.5 ± 1.1 | 6.6 ± 1.3 | 8.5 ± 2.5 | 8.6 ± 0.9 | 133.0 ± 22.8 |
| | FLX-males | 3.9 ± 1.0 | 3.3 ± 1.2 | 4.1 ± 1.0 | 6.4 ± 1.0 | 7.5 ± 1.9 | 7.0 ± 1.2 | 116.3 ± 17.7 |
| All sniffing | CTR-females | 59.0 ± 10.7 | 55.1 ± 7.0 | 49.4 ± 4.9 | 66.5 ± 9.3 | 56.3 ± 5.2 | 48.4 ± 3.6 | 90.1 ± 9.4 |
| | FLX-females | 69.5 ± 6.6 | 68.0 ± 8.6 | 62.8 ± 5.1 | 67.4 ± 5.8 | 66.4 ± 9.2 | 48.1 ± 8.2 | 89.0 ± 6.8 |
| | CTR-males | 47.5 ± 8.6 | 44.0 ± 6.7 | 53.8 ± 8.4 | 50.9 ± 8.0 | 50.3 ± 5.1 | 50.3 ± 8.7 | 37.4 ± 6.0 |
| | FLX-males | 42.0 ± 6.6 | 44.8 ± 8.5 | 46.9 ± 3.9 | 46.9 ± 5.3 | 61.0 ± 6.6 | 51.1 ± 4.9 | 32.1 ± 5.2 |
| Sniffing body | CTR-females | 26.9 ± 5.8 | 28.0 ± 5.6 | 23.6 ± 4.1 | 31.8 ± 6.4 | 27.0 ± 3.8 | 21.6 ± 2.3 | 158.9 ± 24.6 |
| | FLX-females | 35.1 ± 5.3 | 35.6 ± 4.9 | 30.3 ± 4.1 | 31.0 ± 3.2 | 32.6 ± 5.7 | 24.3 ± 5.1 | 188.9 ± 22.8 |
| | CTR-males | 23.6 ± 3.1 | 19.1 ± 1.6 | 21.3 ± 3.5 | 21.0 ± 3.2 | **19.9 ± 2.3** | 21.3 ± 2.8 | 126.1 ± 9.9 |
| | FLX-males | 23.8 ± 4.4 | 20.0 ± 3.1 | 22.0 ± 1.9 | 24.3 ± 3.5 | **30.6 ± 3.3** | 23.6 ± 2.6 | 144.3 ± 14.7 |
| Fighting | CTR-females | 3.8 ± 0.9 | 5.0 ± 1.2 | 4.9 ± 1.2 | 7.8 ± 1.0 | 5.8 ± 1.9 | 5.0 ± 0.8 | 32.1 ± 3.9 |
| | FLX-females | 5.4 ± 1.5 | 6.5 ± 1.6 | 6.5 ± 1.3 | 6.4 ± 1.1 | 4.9 ± 1.1 | 5.0 ± 1.8 | 34.6 ± 5.2 |
| | CTR-males | 0.8 ± 0.4 | 1.4 ± 0.5 | 1.6 ± 0.5 | 2.4 ± 0.7 | 2.3 ± 0.8 | 1.3 ± 0.5 | 9.6 ± 1.9 |
| | FLX-males | 0.9 ± 0.5 | 1.1 ± 0.5 | 2.6 ± 0.5 | 5.1 ± 2.2 | 3.1 ± 0.9 | 1.8 ± 0.7 | 14.6 ± 3.4 |
| Nose-off | CTR-females | 0.0 ± 0.0 | 0.0 ± 0.0 | 0.0 ± 0.0 | 0.0 ± 0.0 | 0.0 ± 0.0 | 0.0 ± 0.0 | 0.0 ± 0.0 |
| | FLX-females | 0.0 ± 0.0 | 0.0 ± 0.0 | 0.0 ± 0.0 | 0.1 ± 0.1 | 0.0 ± 0.0 | 0.0 ± 0.0 | 0.1 ± 0.1 |
| | CTR-males | 0.0 ± 0.0 | 0.0 ± 0.0 | 0.0 ± 0.0 | 0.8 ± 0.5 | 0.4 ± 0.2 | 0.6 ± 0.4 | 1.8 ± 0.9 |
| | FLX-males | 0.0 ± 0.0 | 0.0 ± 0.0 | 0.1 ± 0.1 | 1.1 ± 0.5 | 0.8 ± 0.3 | 0.6 ± 0.4 | 2.5 ± 0.8 |
| Self-grooming | CTR-females | 2.5 ± 0.5 | 3.0 ± 1.1 | 4.5 ± 1.0 | 4.1 ± 1.0 | 5.0 ± 1.4 | 5.5 ± 1.2 | 24.6 ± 3.8 |
| | FLX-females | 2.3 ± 0.7 | 3.3 ± 0.8 | 3.6 ± 0.5 | 3.5 ± 1.0 | 3.0 ± 0.9 | 6.4 ± 1.6 | 22.0 ± 2.5 |
| | CTR-males | 0.4 ± 0.3 | 0.9 ± 0.3 | 0.6 ± 0.4 | 1.4 ± 0.7 | 1.3 ± 0.4 | 2.0 ± 0.6 | 6.5 ± 1.5 |
| | FLX-males | 0.3 ± 0.2 | 0.6 ± 0.3 | 1.9 ± 0.6 | 2.4 ± 1.3 | 1.3 ± 0.5 | 1.5 ± 0.5 | 7.9 ± 1.7 |
| Freezing | CTR-females | 0.0 ± 0.0 | 0.1 ± 0.1 | 0.4 ± 0.3 | 2.6 ± 1.6 | 1.0 ± 0.6 | 0.5 ± 0.3 | 4.6 ± 2.4 |
| | FLX-females | 0.0 ± 0.0 | 0.1 ± 0.1 | 0.6 ± 0.4 | 2.1 ± 1.0 | 1.6 ± 0.5 | 1.0 ± 0.3 | 5.5 ± 1.6 |
| | CTR-males | 0.3 ± 0.2 | 0.0 ± 0.0 | 0.3 ± 0.3 | 1.0 ± 0.5 | 0.3 ± 0.2 | 0.4 ± 0.2 | 2.1 ± 0.7 |
| | FLX-males | 0.1 ± 0.1 | 0.0 ± 0.0 | 0.3 ± 0.3 | 0.8 ± 0.4 | 0.3 ± 0.2 | 1.3 ± 0.4 | 2.6 ± 0.6 |
| Rearing supported | CTR-females | 25.6 ± 5.3 | 32.0 ± 5.0 | 27.1 ± 4.4 | 25.9 ± 4.6 | 29.8 ± 3.3 | 30.1 ± 3.7 | 170.5 ± 17.2 |
| | FLX-females | 34.9 ± 4.7 | 40.1 ± 3.7 | 27.1 ± 2.1 | 27.4 ± 4.8 | 24.8 ± 4.1 | 27.0 ± 3.5 | 181.3 ± 14.6 |
| | CTR-males | 10.3 ± 2.1 | 16.5 ± 2.5 | 12.6 ± 2.4 | 11.9 ± 3.0 | 8.9 ± 1.9 | 9.3 ± 2.6 | 69.5 ± 9.1 |
| | FLX-males | 10.9 ± 2.7 | 16.3 ± 3.2 | 12.4 ± 2.4 | 8.8 ± 2.1 | 8.0 ± 2.0 | 8.0 ± 1.3 | 64.6 ± 7.7 |
| Rearing unsupported | CTR-females | 1.3 ± 0.3 | 1.4 ± 0.6 | 1.6 ± 0.4 | 1.1 ± 0.3 | 2.8 ± 0.8 | 1.5 ± 0.6 | 9.6 ± 1.6 |
| | FLX-females | 0.9 ± 0.3 | 2.3 ± 1.0 | 2.6 ± 2.1 | 2.5 ± 1.2 | 3.4 ± 0.9 | 2.0 ± 0.7 | 13.6 ± 2.8 |
| | CTR-males | 0.5 ± 0.3 | 0.4 ± 0.2 | 2.4 ± 1.1 | 2.8 ± 1.0 | 1.0 ± 0.4 | 2.4 ± 1.1 | 9.4 ± 3.4 |
| | FLX-males | 0.1 ± 0.1 | 1.1 ± 0.7 | 3.3 ± 1.2 | 1.0 ± 0.4 | 1.0 ± 0.4 | 4.9 ± 2.2 | 11.4 ± 2.4 |

*Note.* The data represent the number of instances performing all behaviors measured within the six time-bins. Data are shown in mean ± standard error of the mean.
